# Munc18 binds to and organizes membrane-bound acceptor Q-SNARE complexes in a fashion that depends on the membrane’s lipid composition

**DOI:** 10.64898/2026.07.14.738512

**Authors:** Weronika Tomaka, Mark Kreutzberger, Huan Bao, Volker Kiessling, Lukas K. Tamm

## Abstract

Neuroendocrine cells communicate with other cells by releasing neurotransmitters or hormones by exocytosis, which involves SNARE-mediated fusion between secretory vesicles and the plasma membranes of the secreting cells. In neurons two plasma membrane SNARE proteins, Syntaxin-1a and SNAP25, join with the vesicle membrane SNARE protein Synaptobrevin-2 to form a four-helix bundle, which drives membrane fusion. The assembly of these SNAREs, which is highly orchestrated in cells, has been intensely studied in solution using fragments of the SNARE proteins without their transmembrane domains or lipid anchors. However, in cell and model membranes, Syntaxin and SNAP25 are known to oligomerize and cluster, and little is known about how clustering affects their incorporation into SNARE complexes. In cells, the SM protein Munc18 has been implicated in aiding secretory vesicle docking and facilitating SNARE complex assembly through its interactions with Syntaxin.

To understand how Munc18 orchestrates SNARE complex assembly on membranes, we employed protein reconstitution in model membranes as well as biochemical and biophysical assays to show that lipid-dependent oligomerization of Syntaxin affects Munc18-Syntaxin binding and SNAP25 insertion into the plasma membrane acceptor SNARE complex. We showcase the consequences of the different modes of Munc18-Syntaxin and SNAP25 interaction on Syntaxin’s oligomerization and orientation relative to the membrane surface, as well as on docking and fusion of purified insulin granules. We also determined low-resolution structures by cryoEM in nanodiscs and on the surface of proteoliposomes of membrane-bound assembly states of Munc18/Syntaxin and Munc18/Syntaxin/SNAP25 complexes.

## Introduction

Fusion of secretory vesicles with plasma membranes in neurons and other secretory cells is a key event in cellular communication. Intracellular membrane fusion is controlled by many proteins centered around the core fusion machinery composed of soluble N-ethylmaleimide-sensitive factor (NSF) attachment protein receptor (SNARE) proteins. In neuronal exocytosis the three SNARE proteins, Syntaxin-1a, SNAP25, and Synaptobrevin-2, assemble into a four-helix bundle called the SNARE complex. Assembly of this complex is highly exergonic and drives membrane fusion [1, 2]. Syntaxin and SNAP25 are plasma membrane proteins that contribute one and two SNARE motifs, respectively, while Synaptobrevin is a vesicle membrane protein that contributes one SNARE motif to the final SNARE complex [3]. SNARE motifs consist of a stretch of 60-70 amino acid residues arranged in heptad repeats of hydrophobic residues that form the contacts in the four-helix bundle. The heptad repeats are interrupted in the central layer with either a glutamine (Q) or an arginine (R) instead of a hydrophobic residue. The target membrane SNAREs Syntaxin (Syx) and SNAP25 are Q-SNAREs, while the vesicle membrane SNARE Synaptobrevin (Syb) is an R-SNARE. All SNARE complexes are formed by three Q-SNARE motifs and one R-SNARE motif. Moreover, Syntaxin and Synaptobrevin are anchored in their respective membranes by a helical transmembrane domain that upon fusion extends the SNARE motif helices into the membrane [4], while SNAP25 is a soluble protein that is quadruply palmitoylated in a flexible linker region between its two SNARE motifs. This lipidation anchors SNAP25 to the plasma membrane.

Q- and R-SNARE assembly has been extensively studied in solution with constructs that lack transmembrane domains and lipidations. It is well known that Q-SNAREs alone can form 2:1 (Syx:Syx:SNAP25) complexes [5–8]. The kinetics of SNARE assembly in solution have been studied recently in detail, and many assembly intermediates have been characterized [9]. It was concluded from these studies that the N-terminal SNARE motif of SNAP25 (SN1) assembles with Syx first before the C-terminal motif (SN2) or Syb can integrate into the complex.

In membranes, the situation can be quite different. It is well documented that Q-SNAREs, especially Syntaxin, can self-associate and form clusters in model and cell membranes [10–16]. Therefore, the question arises whether SNARE assembly occurs differently on membranes than in solution and how oligomerization and clustering influence the assembly of membrane-bound SNAREs. A recent study concluded that, like their soluble counterparts, membrane bound Q-SNAREs also assemble in a hierarchical manner, namely that SN1 of SNAP25 assembles with Syntaxin first before SN2 can be incorporated into the Q-SNARE complex [17]. However, unlike in solution, lipid-mediated tethering to the membrane within the soluble linker between SN1 and SN2 facilitated cooperative effects between the two segments and their incorporation into the acceptor Q-SNARE complex [17].

It is well known that in cells Sec1/Munc18-like (SM) proteins and CATCHR proteins like Munc13 are required to dock secretory vesicles to Q-SNARE-rich sites in the plasma membrane and, presumably, to prepare Q-SNAREs for the insertion of the vesicle R-SNARE in preparation for fusion [13, 14, 18–20]. This process is often referred to as “priming” (for fusion) of the SNARE complex although it is not really understood what priming means in mechanistic molecular terms. The Q-SNARE-rich sites, possibly clusters of Q-SNAREs, in the plasma membrane are also known to be rich in phosphoinositides such as PI(4,5)P_2_ and PI(3,4,5)P_3_ [12, 16, 21, 22]. Indeed, the clustering of Syntaxin has been shown in lipid model membranes to depend on PI(4,5)P_2_ and cholesterol [10–12, 23]. PI(4,5)P_2_ binds to a polybasic region between the SNARE motif and transmembrane domain of Syntaxin [11].

It is not clear if the binding of SM and CATCHR proteins to Q-SNAREs and the insertion of the vesicle-bound R-SNARE Synaptobrevin requires a full or partial dissociation of Q-SNAREs from Q-SNARE clusters on the plasma membrane. However, it is known that the SM protein Munc18 binds to Syntaxin in two different modes. In one form Munc18 binds to the so-called closed conformation of Syntaxin in which the Habc domain is folded back onto its SNARE motif, which prevents assembly of Syntaxin with other SNAREs including with itself [24–26]. In another form, Munc18 interacts with the Syntaxin/SNAP25 Q-SNARE complex where Syntaxin’s conformation is slightly open [27, 28]. This form has been proposed to function as an intermediate in SNARE assembly [27–29].

An alternative assembly model based on crystal and cryoEM structures of Munc18 with the soluble SNARE motifs of Syntaxin and Synaptobrevin (or their analogs) [30, 31] posits that Munc18 with further assistance of Munc13 templates the assembly of Syntaxin and Synaptobrevin SNARE motifs in an initial trans-SNARE complex intermediate, into which SNAP25 is subsequently inserted. It is not clear if this complex exists as a defined entity on membranes.

In the present work we designed experiments to illuminate how Munc18 interacts with Q-SNAREs on membranes, which as afore mentioned, could be quite different from analogous interactions that have been observed in solution. Specifically, we wished to address the following questions: How does the oligomerization of Syntaxin on membranes affect the binding of Munc18? How is the binding of Munc18 affected by the lipid composition of the membrane, most notably by acidic lipids and the presence of PI(4,5)P_2_, which we know affects the clustering of Syntaxin? Does Syntaxin oligomerization and Munc18 binding change when Q-SNARE complexes are co-reconstituted with membrane-bound SNAP25? Does docking and fusion of secretory vesicles change when the oligomerization state of the Q-SNAREs and their status of Munc18 binding changes? Since in the course of this work we also found that Munc18 can form relatively stable complexes with Syntaxin alone or with the Syntaxin/SNAP25 Q-SNARE complexes, we also determined low resolution cryoEM structures of these complexes in native nanodiscs that were formed from these membrane-reconstituted complexes. Our low resolution structures confirm that defined Munc18/Syntaxin and Munc18/Syntaxin/SNAP25 complexes exist in membranes and suggest a difference in Munc18 interactions to Syntaxin alone and to Syntaxin in a Syntaxin/SNAP25 Q-SNARE complex. These results set the stage for future higher resolution structural studies.

## Results

### The oligomerization of Syntaxin and its ability to form complexes with SNAP25 on membranes depends on the membrane’s lipid environment

Munc18 has been shown to be necessary for Syntaxin recruitment to the sites of vesicle docking [18, 19]. However, the lipid environment has also been shown to greatly affect Syntaxin’s lateral organization in membranes [10–12, 16, 21–23, 32, 33]. In particular, the formation of Syntaxin clusters strongly depends on its interactions with phosphatidylinositides such as PI(4,5)P_2_ and PI(3,4,5)P_3_ [12, 16, 21, 22]. Therefore, an efficient docking of secretory granules in cells requires a carefully tuned interplay of Munc18-Syntaxin interactions [20] and the formation of Syntaxin clusters in the plasma membrane [13, 14, 23]. Hence, we designed experiments to examine the interdependence of Syntaxin’s lipid environment and Munc18-Syntaxin interactions and how this relationship is further modulated by the presence and interaction with the other Q-SNARE SNAP25. To do this, we took a ground-up approach testing for: (1) the effect that lipid composition has on oligomerization when Syntaxin is alone, (2) the effect that SNAP25 has on Syntaxin oligomerization in those same lipid environments, and (3) how the oligomerization under all these conditions impacts Munc18 binding to Q-SNAREs.

To that end, we reconstituted Syntaxin in membranes of two different lipid compositions. The first was a simple mix of brainPC (bPC) and cholesterol (Chol) (bPC:Chol, 70:30 ratio), which is similar to PC:Chol membranes often used to study SNARE proteins and which lacks any charged lipids. The second lipid composition was a complex mixture that resembles the composition of the plasma membrane (PM: bPC:bPE:bPS:Chol:PI:PI(4,5)P_2_, 25:25:15:30:4:1 ratio). We name this mixture PM 1% PIP2. To test whether the presence of SNAP25 and its potential association with Syntaxin to form an acceptor Q-SNARE complex plays a role in the interdependence of Syntaxin self-association and Munc18 binding, we also used membrane-bound dodecylated SNAP25 (dSNAP25) in these experiments [34]. When dSNAP25 was included, Syntaxin and dSNAP25 were co-reconstituted together in a molar ratio of 1:1 during proteoliposome preparation, which allowed them to form various complexes with each other without forcing any specific stoichiometry.

To probe for the oligomerization of Syntaxin in model membranes we utilized the self-quenching properties of Alexa Fluor 647. Fluorescence self-quenching is very sensitive at close inter-fluorophore distances (<∼1 nm) and since Syntaxin interacts with lipids in its juxta-membrane region [11], the transmembrane region of Syntaxin would respond the strongest to lipid-dependent oligomerization compared to other regions of Syntaxin. We therefore attached Alexa 647 to the C-terminus of Syntaxin (at an additional cysteine in position 289) (Fig. 1A). Consistent with previous reports [10, 11], Syntaxin was more oligomerized in lipid membranes lacking charged lipids, and more dispersed in PM 1% PIP2 membranes also containing PE, PS, PI, and PI(4,5)P_2_ (Fig. 1B). Co-reconstitution with SNAP25 increased the observed fluorescence only slightly in both lipid environments (Fig. 1B), likely because our fluorescence quenching assay probes the oligomerization of the transmembrane domains and not the behavior of the SNARE motifs.

**Figure 1.**
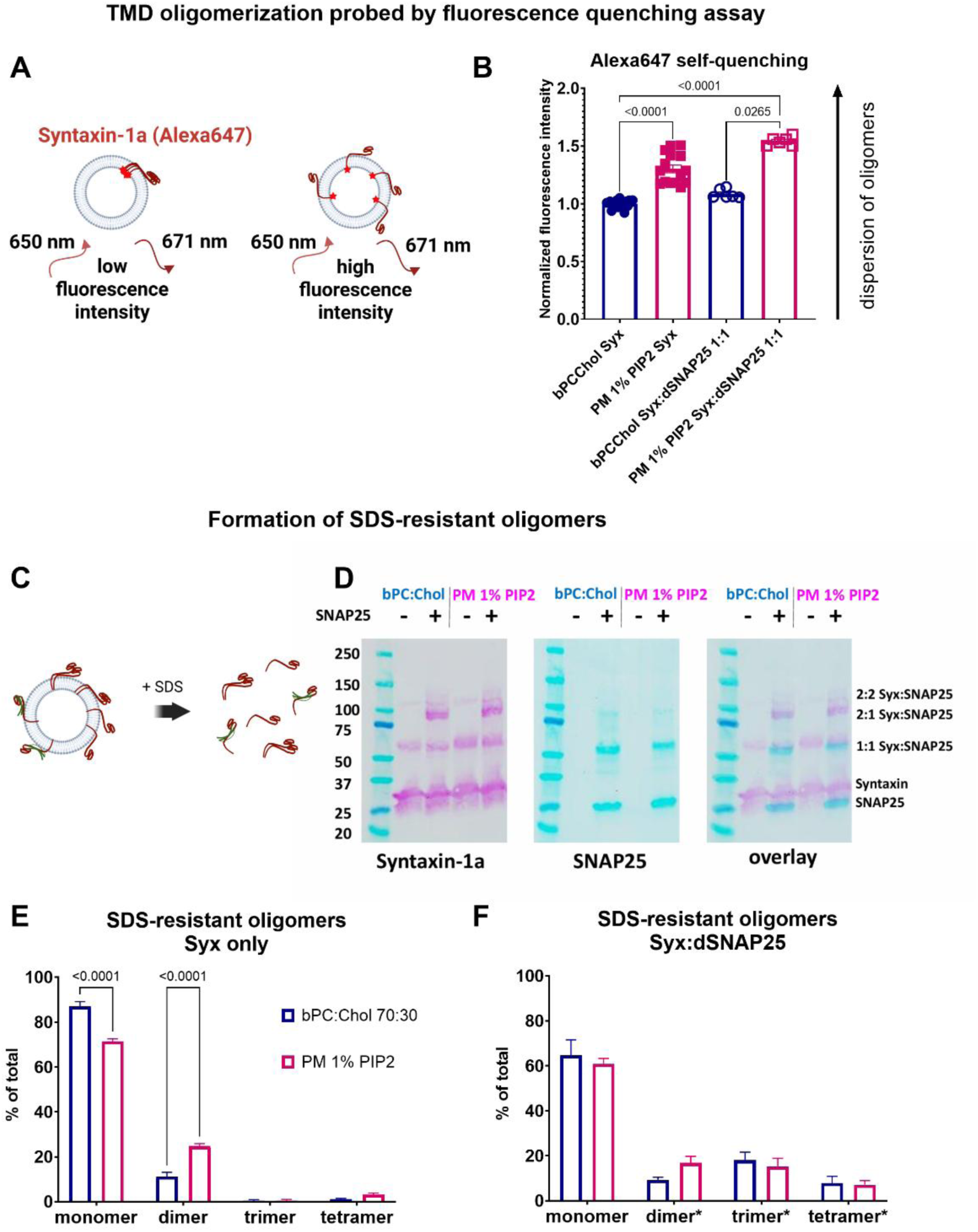
Syntaxin forms complexes with itself and with SNAP25 in a lipid environment dependent manner. **(A)** Self-quenching of Alexa 647 attached to the C-terminus of Syntaxin was used to probe oligomerization of Syntaxin in proteoliposomes. **(B)** Alexa 647 fluorescence intensity normalized to bPC:Chol Syx condition. Kruskal-Wallis test for non-Gaussian distribution was used for statistical analysis. **(C)** Probing the state of Q-SNAREs by solubilizing Q-SNARE proteoliposomes in SDS. **(D)** SDS gel electrophoresis of solubilized Q-SNAREs and Western blotting with anti-Syntaxin-1a (left) and anti-SNAP25 (right) antibodies. **(E and F)** Quantification of populations of oligomeric states of Syntaxin and SNAP25 from Syntaxin only **(E)** and Syntaxin/SNAP25 **(F)** samples. In **(F)** oligomer populations are marked with asterisks to indicate that they might consist of the mixture of Syntaxin’s homo-oligomers and Syntaxin/SNAP25 hetero-oligomers. Two-way ANOVA was used. Error bars represent standard error.

Our fluorescence self-quenching assay is an ensemble measurement that does not allow us to discern what types of complexes are formed. The SNARE complex is SDS-resistant [35, 36], and it has been shown previously that binary Syntaxin-Synaptobrevin complexes are SDS-resistant as well [37]. We utilized this feature to determine the oligomerization state of Syntaxin reconstituted in liposomes under the same experimental conditions that were used in the fluorescence experiments (Fig. 1C). After solubilization in SDS, unboiled samples were run on SDS gels and Western blotted (Fig. 1D). In both tested lipid compositions Syntaxin is mostly monomeric, albeit with significant homo-dimer populations. The Syntaxin dimer population appears to be higher in the charged PM 1% PIP2 than in the uncharged bPC:Chol lipid environment (Fig. 1E). The apparent discrepancy with the fluorescence quenching results may be explained by a model in which charged lipids prevent oligomerization of the transmembrane portion of Syntaxin but still allow oligomerization of Syntaxin’s SNARE motifs [15, 38, 39]. When Syntaxin was co-reconstituted with SNAP25, we observed bands of higher molecular weight that could correspond to 2:1 and 2:2 Syntaxin:SNAP25 complexes (Fig. 1D and F).

Overall, our fluorescence self-quenching and SDS resistance results emphasize the importance of the lipid environment as a prerequisite for regulating Syntaxin’s organization. They explain why Syntaxin’s interaction with charged lipids is crucial in cell membranes [16, 23]. The experiments also show that Syntaxin can form stable complexes with SNAP25 in membranes.

### The binding of Munc18 to Syntaxin and Syntaxin/SNAP25 complexes on membranes depends on the Q-SNARE’s lipid environment

We next explored how the binding of Munc18 to Syntaxin is affected by Syntaxin’s state of oligomerization and interaction with the other Q-SNARE SNAP25 in plasma membrane-mimicking and more simple uncharged lipid bilayers. First, we assessed the binding of fluorescently labeled Munc18 to Syntaxin in planar supported lipid bilayers using TIRF microscopy (Fig. 2A) by recording mean fluorescence intensity of the field of view over time (Fig. 2B and D). Munc18 binds more strongly to Syntaxin in bilayers resembling the plasma membrane (PM 1% PIP2) compared to bilayers composed of only bPC:Chol (Fig. 2B, C). Co-reconstitution of Syntaxin with SNAP25 further increased Munc18 binding (Fig. 2D, E) in both lipid environments. To rule out the possibility that the increased binding in PM 1% PIP2 membranes is due to unspecific interactions of Munc18 with charged lipids or membrane defects, we preincubated the membranes with 2 μM ovalbumin (OVA) before the addition of Munc18 to mask potential non-specific binding sites (Supplementary Fig. 1). After subtracting the residual non-specific binding in the presence of ovalbumin, we again saw that Munc18 still binds more strongly to Syntaxin in PM 1% PIP2 membranes compared to bPC:Chol membranes (Supplementary Fig. 1C, D). Likewise, Munc18 binding is enhanced in the presence of SNAP25 even after pre-saturating the membranes with ovalbumin (Supplementary Fig. 1E, F).

**Figure 2.**
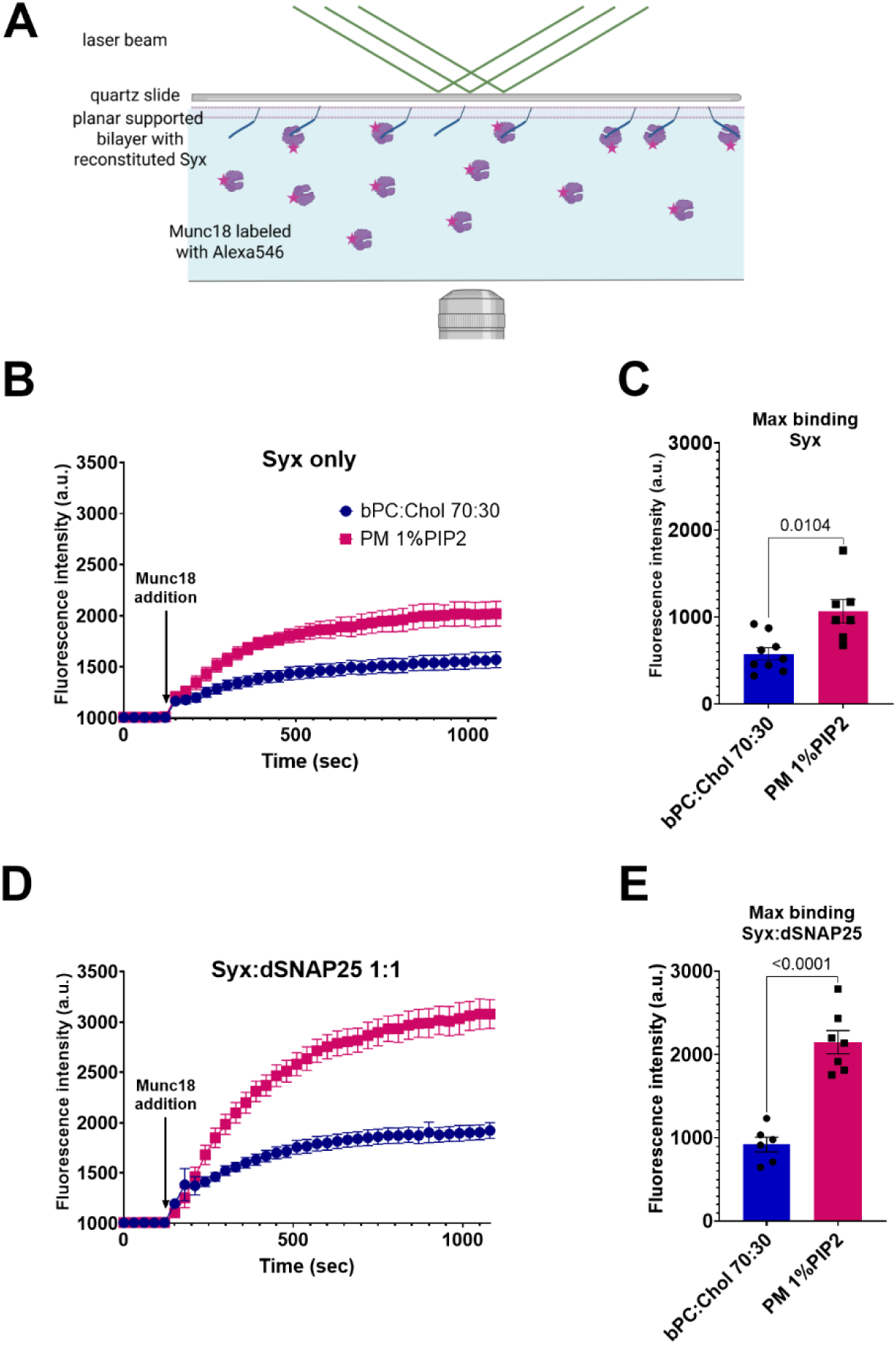
Charged lipids and co-reconstitution with SNAP25 increase Munc18 binding to Syntaxin in planar supported membranes. **(A)** TIRF microscopy was used to examine binding of fluorescently labeled Munc18 to planar supported membranes. **(B)** TIRF microscopy binding curves of 0.5 µM Munc18 to planar supported membranes containing Syntaxin only. **(C)** Maximum fluorescence intensity values obtained from fits of the binding curves from **(B)** to 1^st^ order kinetic functions. **(D)** TIRF microscopy binding curves of 0.5 µM Munc18 to planar supported membranes containing Syntaxin co-reconstituted with SNAP25. **(E)** Maximum fluorescence intensity values obtained from fits of the binding curves from **(D)** to 1^st^ order kinetic functions. Error bars represent standard error. T-test with Welch’s correction for unequal SDs was used in **(C)** and **(E)**. We additionally used t-test with Welch’s correction to compare maximum fluorescence intensity values between Syntaxin only and Syntaxin:SNAP25 membranes of each lipid condition. Differences were statistically significant, with p=0.0001 for PM 1% PIP2 membranes, and p=0.0113 for bPC:Chol membranes.

To achieve additional insight into the interaction of Munc18 with Syntaxin on membranes, we performed co-floatation assays (Fig. 3A). In this approach, Munc18 is allowed to bind to Syntaxin reconstituted into liposomes of different lipid compositions with or without SNAP25 before the samples are deposited with 40% Nycodenz at the bottom of a Nycodenz density gradient and spun at 2x10^5^ g for 1.5 hours. Free protein sediments at the bottom and proteoliposomes with any bound protein float up to the top of the gradient in this assay. Fractions are analyzed by fluorescence for lipid or Western blotting for proteins. A lipid marker and Syntaxin follow a smooth distribution in the gradient with Syntaxin trailing the lipids slightly in PM 1% PIP2 membranes when Munc18 is bound (Fig. 3B, C). With bPC:Chol Syntaxin proteoliposomes, a large fraction of Munc18 migrates together with Syntaxin to the top fractions of the gradient, suggesting formation of stable Munc18/Syntaxin complexes (Fig. 3D). However, with PM 1% PIP2 Syntaxin proteoliposomes, the distribution of Munc18 is shifted towards fractions 2, 3, and 4 (of 10), which could be an indication of a weaker, more easily dissociating Munc18/Syntaxin complex on PM 1% PIP2 membranes in this non-equilibrium experiment (Fig. 3D). In both lipid conditions, about 50% of Munc18 remains unbound in the bottom fractions (7, 8, 9, and 10). Interestingly, when Syntaxin is co-reconstituted in proteoliposomes with SNAP25 prior to Munc18 binding, slightly less Munc18 partitions to top fractions in PM 1% PIP2 membranes and none in bPC:Chol membranes (Fig 3E). Since we observed more Munc18-Syntaxin association in planar supported membranes in presence of SNAP25 compared to Syntaxin alone (Fig. 2), this result suggests that Munc18 interacts readily with the Syntaxin/SNAP25 complex, but that the interaction with this complex may form a less stable intermediate than with Syntaxin alone.

**Figure 3.**
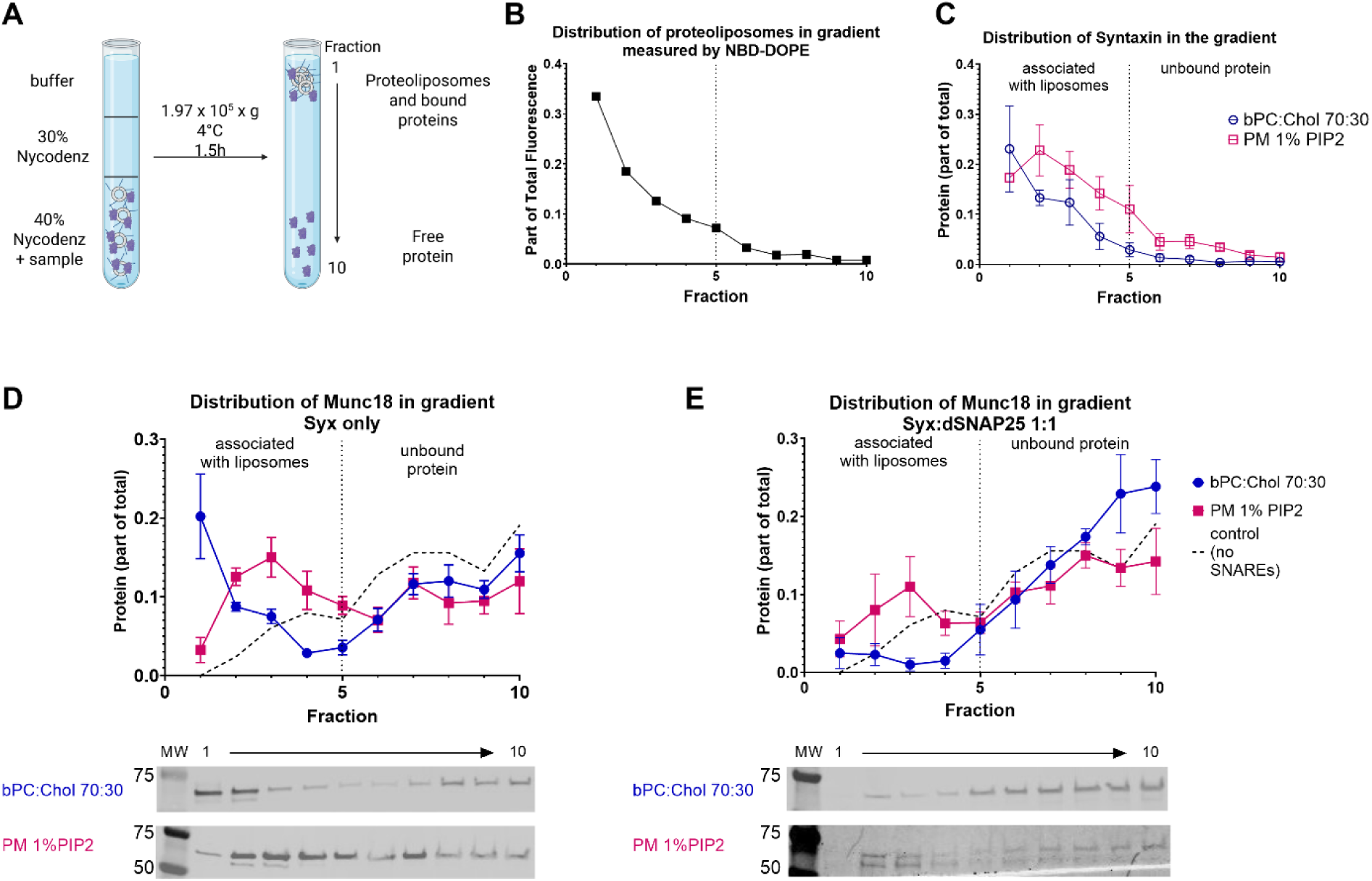
Munc18 distribution in co-floatation assay with Q-SNARE proteoliposomes reveals differences in the stability of binding of Munc18 to Syntaxin only and to Syntaxin/SNAP25 complexes. **(A)** Design of the co-floatation assay in a Nycodenz density gradient. **(B)** Distribution of lipids probed by NBD-DOPE in PM proteoliposomes in the gradient. **(C)** Distribution of Syntaxin in PM and bPC:Chol proteoliposomes in the gradient. **(D, E)** Distribution of Munc18 in the gradient with PM and bPC:Chol proteoliposomes with Syntaxin only **(D)** and Syntaxin co-reconstituted with SNAP25 **(E)**. Error bars indicate standard error of the mean.

Taken together, these results suggest that the acidic lipids (i.e., 1 mol% PIP2, 4 mol% PI, and 15 mol % PS) and 25 mol% PE in the PM 1% PIP2 mix decrease the tight packing of Syntaxin’s transmembrane domains, while simultaneously allowing for increased small-scale oligomerization of Syntaxin’s SNARE motif in PM 1% PIP2 vs. bPC:Chol membranes (Fig. 1). Combination of these two effects results in higher accessibility of Syntaxin for interactions with Munc18 (Fig. 2). Increased binding of Munc18 persists or is even enhanced when Syntaxin is co-reconstituted with SNAP25 (Fig. 2) and its small-scale oligomerization is increased (Fig. 1), but at the same time Munc18 complexes with Syntaxin/SNAP25 Q-SNARE complex seem more labile than Munc18 complexes with Syntaxin alone (Fig. 3). These results also suggest that Munc18 binds differently to Syntaxin alone compared to Syntaxin/SNAP25 complexes, which is consistent with previous electron paramagnetic resonance (EPR) and double electron electron resonance (DEER) results on the binding of Munc18 to soluble full-length Syntaxin (without TM domain) and SNAP25 (without lipid anchors) [26]. In both cases, binding to Syntaxin alone results in more stable complexes while binding to Syntaxin/SNAP25 is more transient.

### Munc18 binding modifies the lateral organization of Q-SNAREs in membranes in a lipid-dependent manner

One of the key functions of Munc18 is proposed to be maintaining Syntaxin in a closed conformation to reduce its ability to self-oligomerize [25, 26, 29]. Utilizing the fluorescence quenching and formation of SDS-resistant oligomer assays described above, we observed very minor effects of Munc18 binding on Syntaxin oligomerization (Fig. 4). Even so, it is interesting to observe that Munc18’s effect on Syntaxin oligomerization probed by fluorescence quenching of Alexa647 at the C-terminus of Syntaxin again depended on the lipid environment (Fig. 4A and D). In the absence of charged lipids Munc18 caused a very small fluorescence increase when Syntaxin was reconstituted alone, indicating a possible slight dispersion of Syntaxin oligomers (Fig. 4A). In the presence of SNAP25 the fluorescence was unchanged by Munc18 addition (Fig. 4A). However, in PM 1% PIP2 lipids Munc18 seemed to promote oligomerization of Syntaxin, and this effect was still present, albeit less pronounced, in presence of SNAP25 (Fig. 4D). A control, in which fluorescence in both lipid conditions was measured before and after buffer addition (without adding Munc18) showed only a minimal reduction in fluorescence intensity, likely due to photobleaching (last columns in Fig. 4A and D).

**Figure 4.**
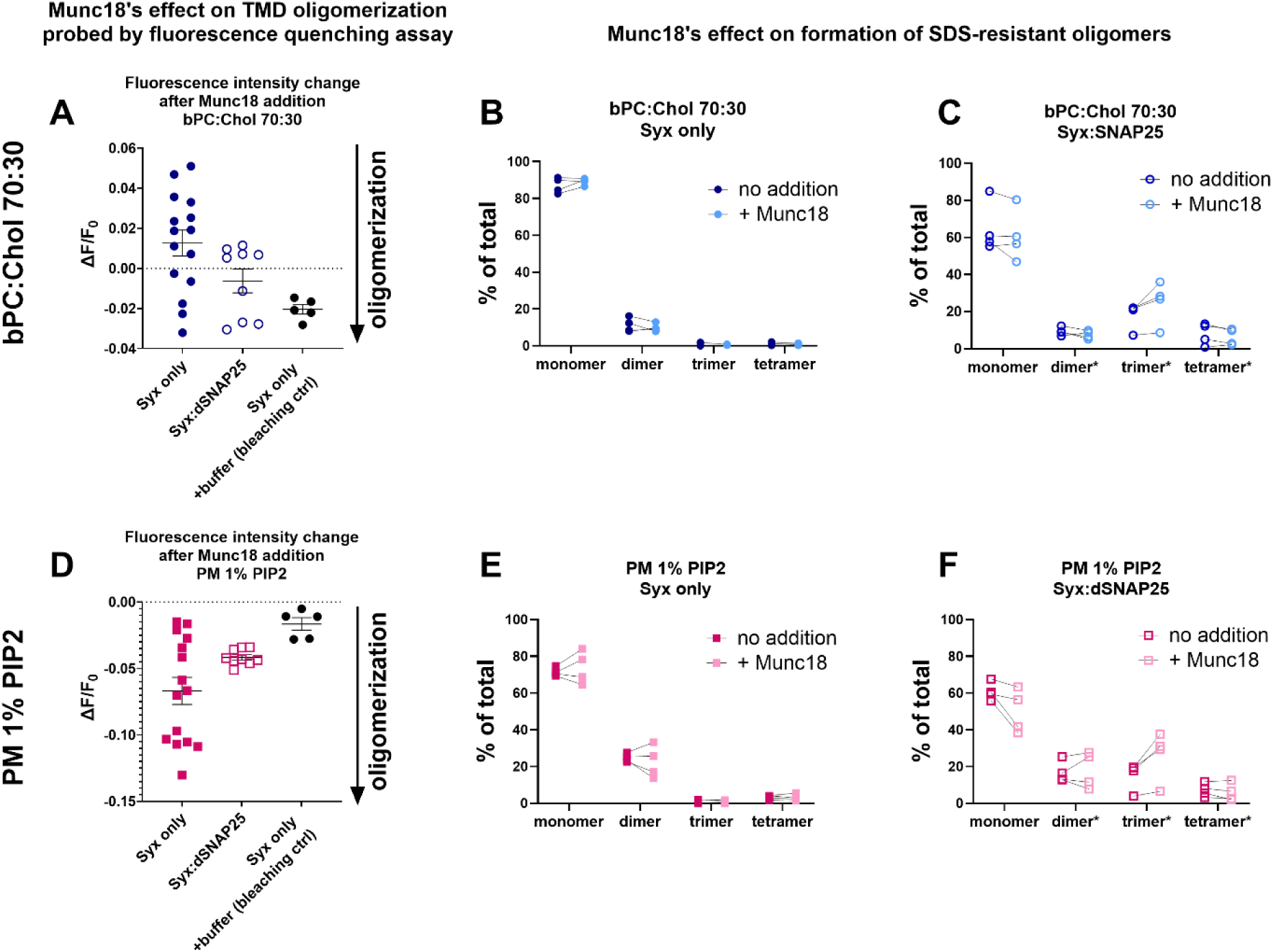
Munc18 binding has only very minor effects on Syntaxin’s oligomerization in bPC:Chol and PM lipid bilayers. **(A)** Change of fluorescence intensity of Syntaxin-289C-Alexa647 in bPC:Chol membranes after the addition of 0.5 µM Munc18. **(B and C)** Quantification of populations of oligomeric states of Syntaxin and SNAP25 from Syntaxin only **(B)** and Syntaxin/SNAP25 **(C)** samples in bPC:Chol membranes before and after incubation with 0.5 µM Munc18. **(D)** Change of fluorescence intensity of Syntaxin-289C-Alexa647 in PM membranes after the addition of 0.5 µM Munc18. **(E and F)** Quantification of populations of oligomeric states of Syntaxin and SNAP25 from Syntaxin only **(E)** and Syntaxin/SNAP25 **(F)** samples in PM membranes before and after incubation with 0.5 µM Munc18.

Examining the SDS-resistant oligomers, we did not notice any change in oligomer population caused by Munc18 when Syntaxin was reconstituted alone in either lipid condition (Fig. 4B and E). Interestingly, in presence of SNAP25 in both lipid conditions Munc18 seemed to slightly decrease the monomer population and promote formation of a trimer (Fig. 4C and F). While the effect is not strong, it is reproducible and suggests that Munc18 might promote formation of Syntaxin/SNAP25 complexes.

Overall, while the observed effects of Munc18 on Syntaxin oligomerization were small, they may point to more nuanced changes that we were unable to observe. Fluorescence self-quenching is only sensitive at small distances and detects only tight interactions between Syntaxin’s trans-membrane domains. We cannot exclude the possibility that Munc18 organizes Syntaxin on a larger scale where Syntaxin packing is not dense enough to be detected by measuring fluorescence self-quenching. It is also possible that the SDS solubilization experiments reveal only an underlying trend of Q-SNARE association and SDS may disrupt some species of higher-order complexes that may be present in lipid bilayers.

Maintaining Syntaxin in its monomeric form is connected to it adopting a closed conformation when bound to Munc18. It has been shown in solution and in PC12 cell membranes that Syntaxin’s N-terminal peptide (residues 1-27) is necessary for Munc18 to bind Syntaxin in a closed conformation [27, 40]. We tested the effect of Munc18 on Syntaxin lacking the N-peptide in the fluorescence self-quenching experiment (Supplementary Fig. 2). In bPC:Chol membranes, the effect of Munc18 on Syntaxin oligomerization was reduced in the absence of the N-peptide of Syntaxin (Supplementary Figure 2B). This effect was not due to a lack of Munc18-Syntaxin binding (Supplementary Fig. 2D and E). In PM 1% PIP2 lipid bilayers, Munc18 seemed to promote Syntaxin oligomerization even stronger for the construct lacking the N-peptide compared to the full-length protein (Supplementary Figure 2C). Taken together, these results suggest that Munc18’s interaction with Syntaxin’s N-peptide is partially responsible for its effect on Syntaxin’s oligomerization in membranes, likely by preventing Munc18-induced closing of Syntaxin’s conformation in the absence of the N-peptide.

### Munc18 binding lifts Syntaxin and Q-SNARE complexes from the membrane surface

The SNARE motif of Syntaxin is connected to its transmembrane domain via a flexible and unstructured linker [41], which becomes alpha-helical as the SNARE complex transitions from the *trans*- to *cis*- configuration in the final stages of membrane fusion [4]. Site-directed Fluorescence Interference Contrast (sdFLIC) microscopy allows to measure ensemble average distances of fluorescent probes from membrane surfaces with sub-nanometer precision [42]. We previously used this technique with multiple labels that were engineered (site-directed) into the SNARE motif of Syntaxin to measure their positions – and therefore the angle of the SNARE motif – from the membrane surface [41, 43]. Here, we utilized sdFLIC microscopy to explore how Munc18 influences Syntaxin’s orientation relative to the membrane surface. To this end, we engineered 7 cysteines along the H_abc_ domain and the SNARE motif of Syntaxin (Fig. 5A), labeled them with Alexa546, and measured their distances from the surface of a reconstituted PM 1% PIP2 supported membrane. In the absence of other proteins, Syntaxin’s SNARE motif and H_abc_ domain are positioned close to the membrane surface (Fig. 5B). The disposition of Syntaxin’s H_abc_ domain does not change much when Syntaxin is co-reconstituted with membrane-anchored SNAP25 and Syntaxin’s SNARE motif distance increases by ∼1 nm from the membrane surface when complexed with SNAP25 (Fig. 5C). However, after Munc18 is bound, the distances of the Habc domain and the SNARE motif are lifted up from the membrane surface by 3-4 nm and the SNARE increases its angle with the membrane surface (Fig. 5B). The same vertical and angular changes of Syntaxin upon Munc18 binding were observed when Syntaxin was co-reconstituted with SNAP25 (Fig. 5C).

**Figure 5.**
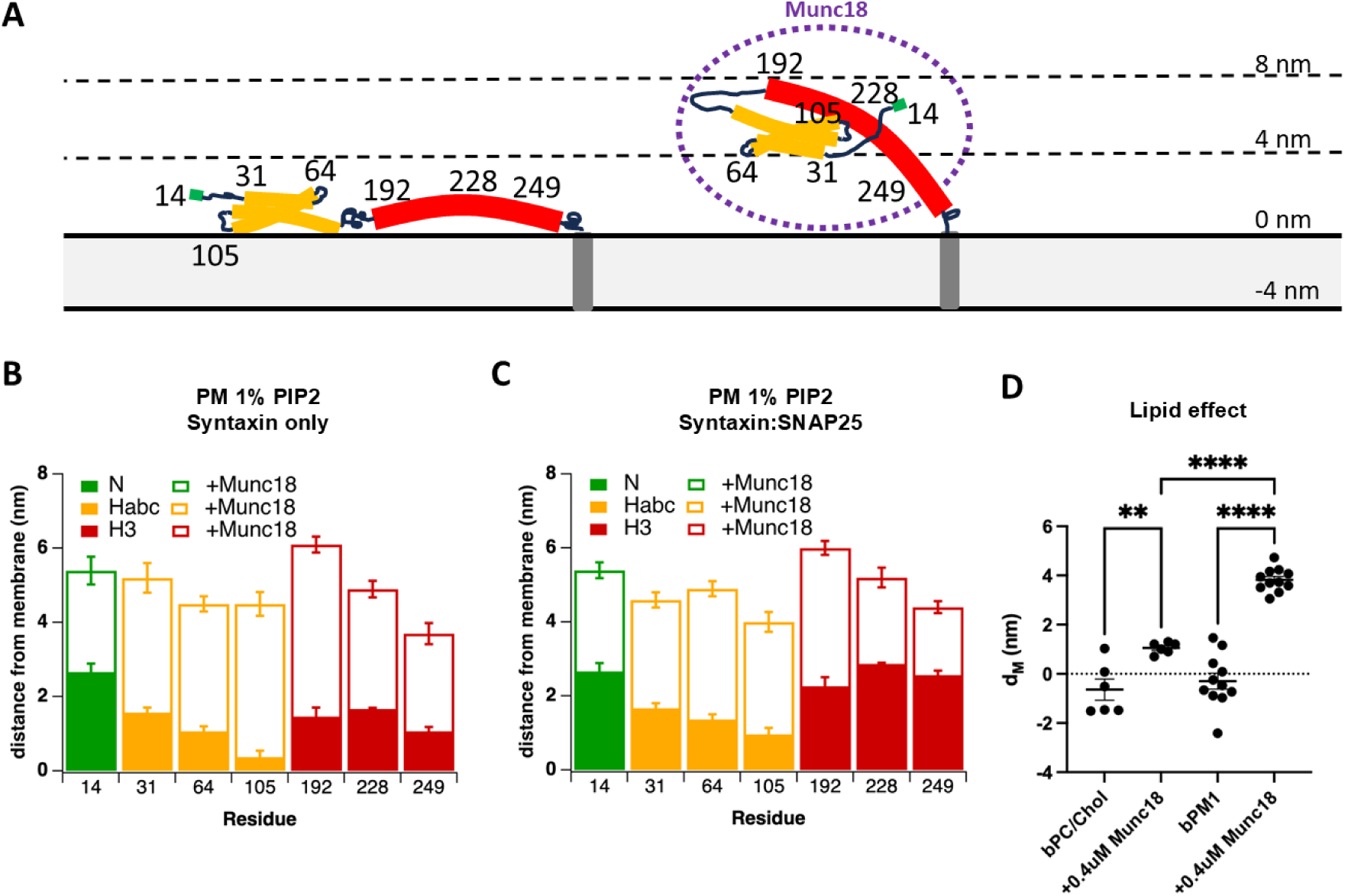
Munc18 elevates Syntaxin from the membrane surface. **(A)** Visual representation of residues of Syntaxin that were site-specifically labeled with Alexa 546 for FLIC microscopy experiments. The N-peptide is marked green, the H_abc_ domain is marked orange, the SNARE motif is marked red, and the 7 labeled residues are indicated. **(B, C)** Distances of the labeled sites of Syntaxin from the membrane before (solid bars) and after (open bars) incubation with 0.5 µM Munc18 in PM 1% PIP2 supported lipid bilayers with Syntaxin only **(B)** or Syntaxin co-reconstituted with SNAP25 **(C)**. **(D)** Distance of residue 192 of Syntaxin (N-terminal end of SNARE motif) co-reconstituted with SNAP25 in bPC:Chol and PM 1% PIP2 lipid bilayers before and after incubation with 0.4 µM Munc18. Error bars represent standard error. For statistical analysis, one-way ANOVA test was used, asterisks indicate ** p=0.003 and **** p<0.0001.

We also examined if the lipid environment had an impact on Munc18’s ability to elevate Syntaxin from the membrane surface. To this end, we measured the FLIC distances of Syntaxin at residue 192 in the Syntaxin/SNAP25 acceptor complex from the surface of a bPC:Chol membrane and compared them with those on PM 1% PIP2 membranes. We found that Munc18 elevates Syntaxin more in PM membranes compared to bPC:Chol membranes (Fig. 5D). Since Munc18 interacts with Syntaxin/SNAP25 more efficiently, albeit in a different mode, on PM 1% PIP2 vs. bPC:Chol membranes (Fig. 2 and 3), these differences in binding affinity and mode likely explain the differences in how efficiently Munc18 changed Syntaxin’s orientation on the membrane surface.

### Syntaxin/SNAP25’s lipid environment and interaction with Munc18 impact docking and fusion of insulin granules with Q-SNARE target membranes

Formation of the Munc18/Syntaxin complex is generally considered to be a crucial step in the assembly of the SNARE complex [2]. The complex likely plays a significant role as a starting point of the assembly and also as a facilitator for incorporating other SNAREs into the nascent complex [30, 31, 44–50]. Since we observed that Syntaxin’s lipid environment affects how Munc18 interacts with Syntaxin and other SNARE proteins, we reasoned that it would also affect docking and fusion of secretory vesicles.

We measured fusion of insulin granules purified from INS1 cells [51] with Q-SNARE proteoliposomes reconstituted from the same components and in the same fashion as in the other experiments of this study in a bulk liposome fusion assay (Fig. 6A). Insulin granules fused much more efficiently with Syntaxin/SNAP25 reconstituted in PM 1% PIP2 than in bPC:Chol proteoliposomes (Fig. 6B). Preincubating the Q-SNARE proteoliposomes with 0.5 μM Munc18 had no effect on fusion in this assay. Since in this study we wanted to probe the effect of Munc18 on the early intermediates of the SNARE complex assembly, we did not include other regulators such as Munc13, complexin, CAPS, or Ca^2+^ in these experiments that are necessary for highly efficient fusion of purified granules [52, 53].

**Figure 6.**
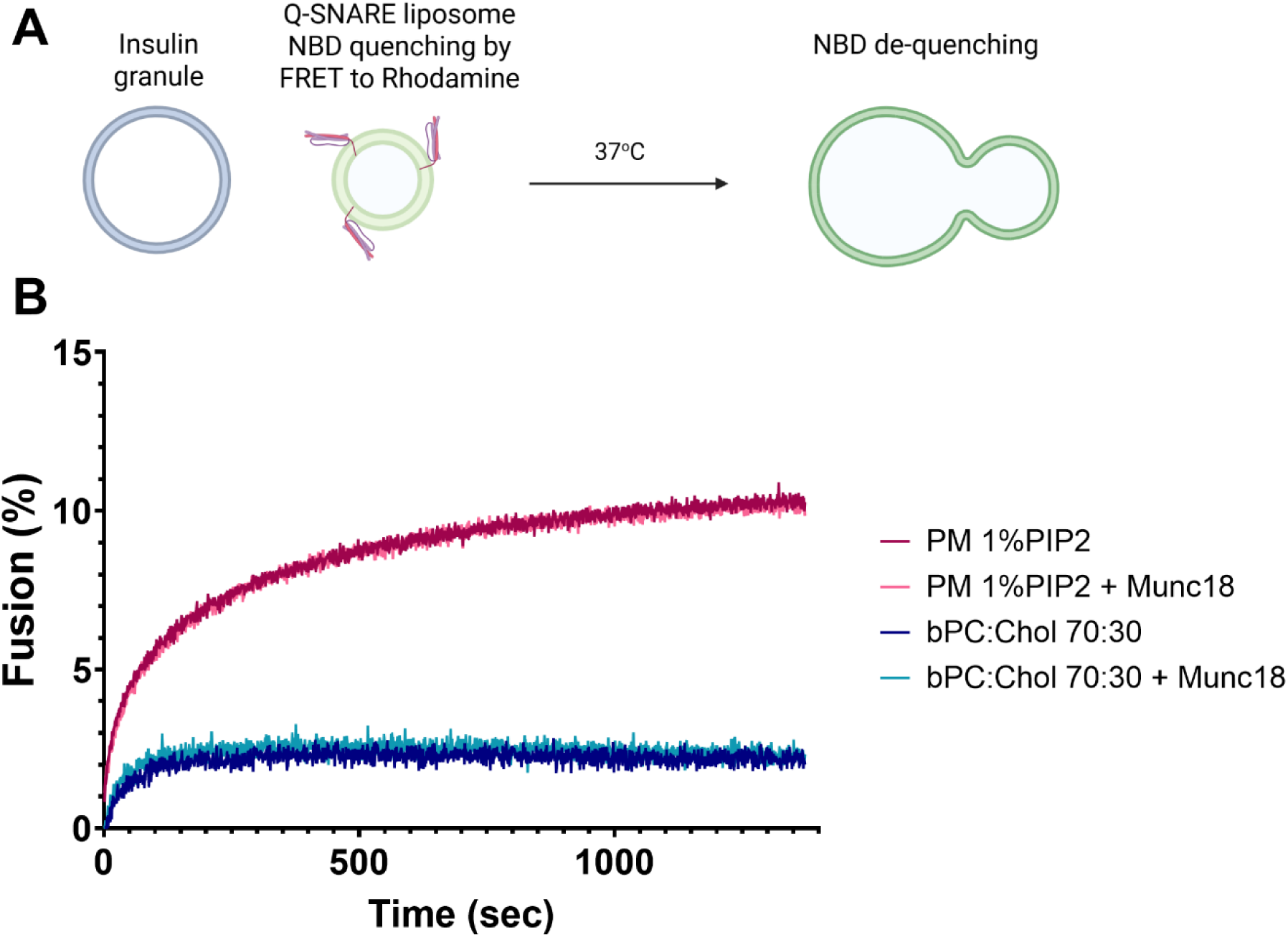
Fusion of insulin granules with Q-SNARE proteoliposomes reveals a strong dependency on lipid environment, but not on the presence or absence of Munc18. **(A)** Q-SNARE proteoliposomes containing the FRET pair NBD-DOPE and Rhodamine-DOPE were mixed with insulin granules isolated from INS1 cells to measure fusion in a cuvette-based bulk fusion assay. **(B)** Fusion of insulin granules with Q-SNARE proteoliposomes with PM 1% PIP2 and bPC:Chol lipid compositions with and without pre-incubation with 0.5 µM Munc18. Percent fusion was determined by NBD dequenching normalized to the total fluorescence after addition of Triton X-100.

The bulk fusion assay reports the convolution of granule docking and fusion efficiencies. To get a detailed picture of the reaction at the single granule level, we imaged binding and fusion of individual insulin granules containing C-peptide-GFP with planar supported membranes using TIRF microscopy [51] (Fig. 7A). An example of a single granule docking and fusion trace is shown in Fig. 7B. Granules docked more efficiently when the Q-SNAREs were reconstituted into PM 1% PIP2 membranes compared to bPC:Chol membranes, likely due to more available Syntaxin in the presence of charged lipids. Munc18 trended to decrease docking slightly with PM 1% PIP2 membranes and more significantly with bPC:Chol membranes, although this may not be statistically significant (Fig. 7C). However, the fusion probability (measured as the percentage of docked granules that undergo fusion) was similar in both lipid conditions and trended to increase when Munc18 was included (Fig. 7D). These opposite effects of Munc18 on docking and fusion likely explain why we didn’t observe Munc18 dependent differences in the bulk fusion assay (Fig. 6).

**Figure 7.**
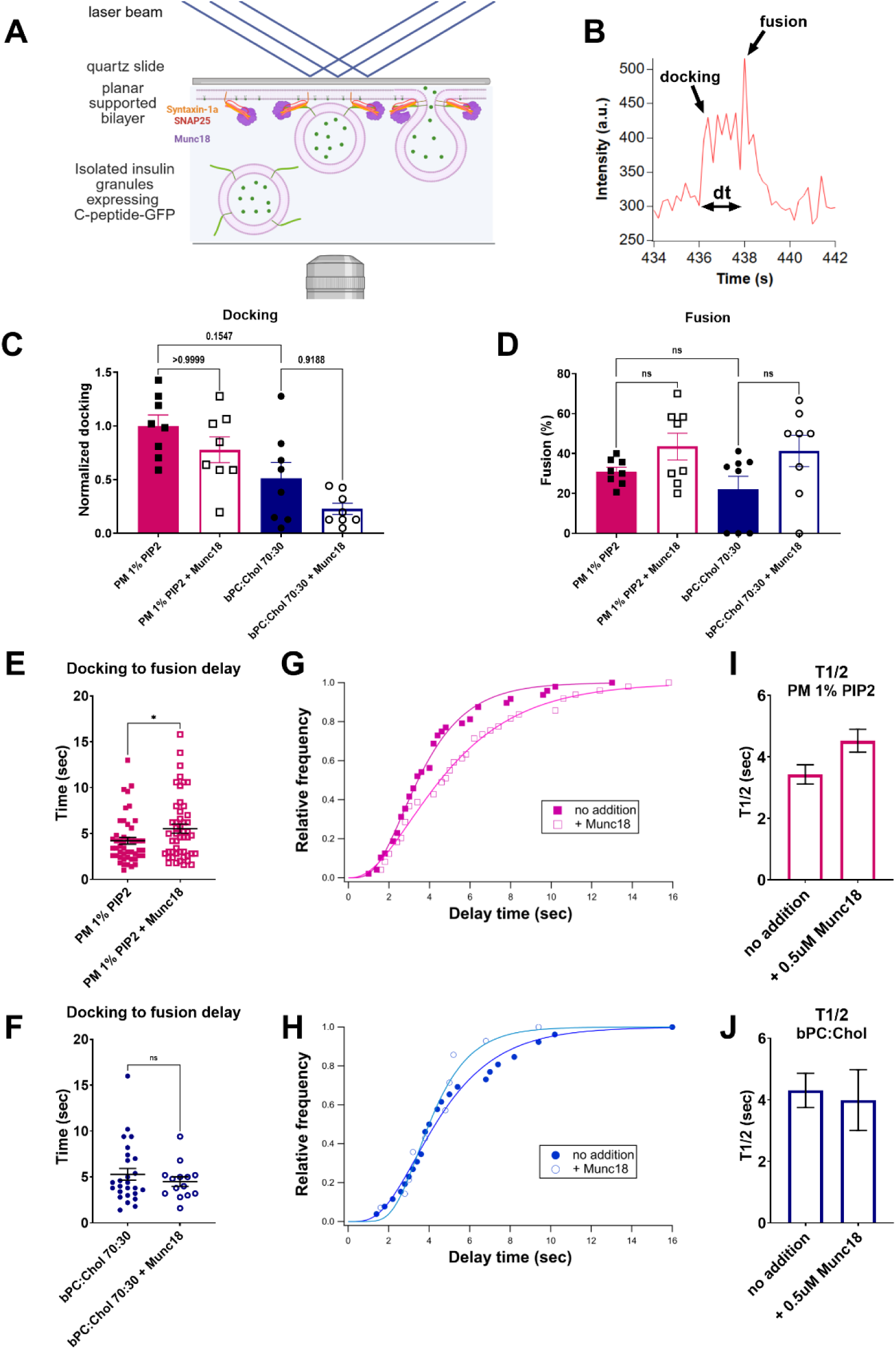
Munc18 affects docking, fusion, and fusion kinetics of insulin granules with planar supported membranes. **(A)** Docking and fusion of insulin granules containing C-peptide-GFP with Q-SNARE-containing planar supported membranes were imaged using TIRF microscopy. **(B)** Example of a fluorescence intensity trace with marked onsets of docking and fusion, and the docking to fusion delay time dt. **(C)** Docking of granules to supported PM 1% PIP2 or bPC:Chol Syntaxin/SNAP25 membranes without and with Munc18, normalized to the number of docked granules with PM 1% PIP2 membranes without Munc18. **(D)** Fusion efficiency measured as a percentage of docked granules that proceeded to fusion under the same conditions as in **(C)**. **(E-J)** Analysis of fusion kinetics. **(E and F)** Comparison of docking to fusion delay times with and without the addition of Munc18 in PM 1% PIP2 membranes **(E)** and bPC:Chol membranes **(F)**. **(G and H)** Cumulative distribution function of the delay times (data points) and the fit with a parallel reaction model *N*(*t*) = (1 − *exp*^−*kt*^)*^m^* [69], where k is the rate constant and m is the number of parallel reactions occurring (solid line) in PM 1% PIP2 **(G)** and bPC:Chol **(H)** membranes. **(I and J)** Half-times of the maximum fusion calculated from the fits of the cumulative distribution function of the delay times in PM 1% PIP2 **(I)** and bPC:Chol membranes **(J)**. For statistical analysis, Kruskal-Wallis test for non-Gaussian distribution was used in multiple comparison tests. Kolmogorov-Smirnov test for cumulative distributions was used to compare docking to fusion delay times.

We also examined the fusion kinetics of the single granule fusion data by analyzing the delay times between granule docking and the onset of fusion (Fig. 7B). Munc18 increased the delay times dt in PM 1% PIP2 membranes (Fig. 7E) but had no effect on the delay times in bPC:Chol membranes (Fig. 7F). The normalized cumulative distribution of the delay times were fitted with a parallel reaction model (see Experimental Procedures) (Fig. 7G and H), which yielded characteristic half-times of maximum fusion for each condition (Fig. 7I and J). These results show that fusion is faster in PM 1% PIP2 membranes (Fig. 7G and I) than in bPC:Chol membranes (Fig. 7H and J), and that Munc18 delays fusion in PM 1% PIP2 membranes (Fig. 7G and I), but not in bPC:Chol membranes (Fig. 7H and J).

Taken together, these results indicate that lipid-dependent oligomerization of Syntaxin affects its availability for docking of secretory granules. Syntaxin is more dispersed in PM 1% PIP2 membranes compared to bPC:Chol membranes, resulting in more docking-competent complexes. Addition of Munc18 to Syntaxin/SNAP25 complexes in bPC:Chol membranes reduces the number of docking-competent complexes available but does not affect the nature of the fusion events (kinetics), which suggests that in both cases (with and without Munc18) fusion is likely mediated by Syntaxin/SNAP25 complexes. In contrast, in PM 1% PIP2 membranes Munc18 only slightly reduces the number of docking-competent sites, but the fusion kinetics are slower, suggesting that fusion now occurs through Munc18/Syntaxin/SNAP25 complexes.

### Cryo-EM of Munc18/Syntaxin and Munc18/Syntaxin/SNAP25 complexes

Throughout this work, we saw evidence for the formation of an acceptor complex consisting of Munc18, Syntaxin and SNAP25. We observed that the Munc18-Syntaxin interaction was affected by presence of SNAP25 (Figs. 2 and 3), and that Munc18 slightly promoted the formation of Syntaxin/SNAP25 complexes (Fig. 4). Our single granule fusion assay further revealed that Munc18 affected the fusion kinetics of granules to plasma membrane-mimicking membranes, leading us to hypothesize that under this condition fusion occurs through a Munc18/Syntaxin/SNAP25 complex-mediated pathway.

We therefore sought to visualize Munc18/Syntaxin and Munc18/Syntaxin/SNAP25 complexes in model membranes using cryo-EM (Fig. 8). We imaged Munc18 bound to Syntaxin/SNAP25 complexes in PM 1% PIP2 proteoliposomes (Fig. 8A-C), and Munc18 bound to Syntaxin and Syntaxin/SNAP25 complexes in nanodiscs derived from the reconstituted PM 1% PIP2 proteoliposomes (Fig 8D-I). The nanodiscs were prepared with the membrane-solubilizing peptide Hex18A, which does not require any detergent for gentle membrane extraction [54] and which therefore preserves even weak membrane protein complexes.

**Figure 8.**
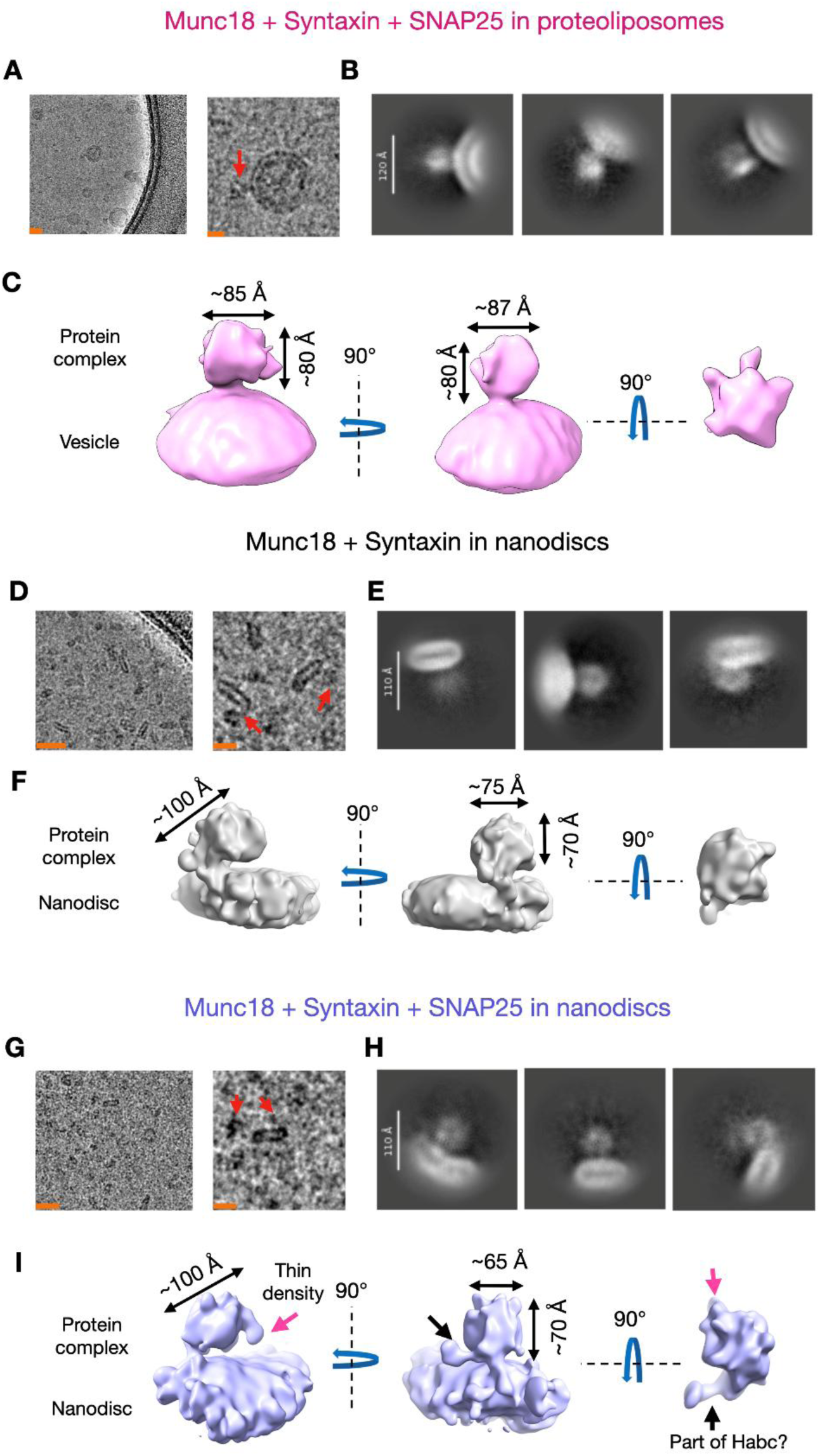
Low resolution cryo-EM structural analysis of Munc18/Syntaxin and Munc18/Syntaxin/SNAP25 complexes. **(A)** Cryo-EM micrographs of PM 1% PIP2 proteoliposomes reconstituted with Munc18/Syntaxin/SNAP25. Scale bars for the images are approximately 200 Å (left) and 100 Å (right). **(B)** 2D class averages of Munc18/Syntaxin/SNAP25 in proteoliposomes. The scale bar is 120 Å. **(C)** Cryo-EM reconstruction of Munc18/Syntaxin/SNAP25 complex in proteoliposomes. **(D)** Cryo-EM micrographs of nanodiscs containing Munc18/Syntaxin complexes. Scale bars for the images are approximately 200 Å (left) and 100 Å (right). **(E)** 2D class averages of Munc18/Syntaxin in nanodiscs. The scale bar is 110 Å. **(F)** Cryo-EM reconstruction of protein complex from Munc18/Syntaxin in nanodiscs. **(G)** Cryo-EM micrographs of nanodiscs containing Munc18/Syntaxin/SNAP25 complexes. Scale bars for the images are approximately 200 Å (left) and 100 Å (right). **(H)** 2D class averages of Munc18/Syntaxin/SNAP25 in nanodiscs. The scale bar is 110 Å. **(I)** Cryo-EM reconstruction of protein complex from Munc18/Syntaxin/SNAP25 in nanodiscs.

From manual picking observations and 2D class averages it is apparent that the protein densities on the surface of the nanodiscs and proteoliposomes were quite heterogenous, which resulted in 2D classes with blurry protein densities for Munc18/Syntaxin and Munc18/Syntaxin/SNAP25 complexes (Fig. 8B, E, H; Supplementary Fig. 3A-C). Besides larger complexes at the surface of the membrane which most likely correspond to Munc18 bound to Q-SNAREs (Fig. 8; Supplementary Fig. 3C), we also observed densities that look like Syntaxin molecules (Supplementary Fig. 3B). Despite extensive particle picking (see Experimental Procedures) we were unable to obtain 2D classes with more defined protein densities. We believe this is likely due to sample heterogeneity and potential flexibility of the complexes. This heterogeneity is consistent with our oligomerization data.

Regardless, we were able to reconstruct low resolution 3D structures for the protein densities of the Munc18/Syntaxin/SNAP25 complex in PM 1% PIP2 proteoliposomes (19 Å resolution) (Fig. 8C), as well as Munc18/Syntaxin (13 Å resolution) and Munc18/Syntaxin/SNAP25 (12.6 Å resolution) complexes in nanodiscs (Fig. 8F and I). Based on the size of the protein densities we believe that each volume corresponded to one Munc18 protein bound to either one Syntaxin or one Syntaxin/SNAP25 complex.

The cryoEM structures of Munc18/Syntaxin and Munc18/Syntaxin/SNAP25 complexes in nanodiscs shared similarities showing globular heads that were connected by necks to the nanodiscs (Fig. 8F and I). Despite these similarities there were also two distinct differences. First, near the neck connection to the membrane there was a large density extending perpendicular to the rest of the complex (Figure 8I and Fig. 9 black arrows). Second, the bilayer connection (neck) of the Munc18/Syntaxin/SNAP25 was narrower than that of the Munc18/Syntaxin complex (Fig. 9) and had an additional thin density on the opposite side of the complex from the neck (Fig. 8I left, Fig. 9, pink arrows). Two half maps of the Munc18/Syntaxin/SNAP25 complex also showed these striking features indicating that the differences were reproducible between the two reconstructions (Supplementary Fig. 4). Additionally, at low density thresholds for the Munc18/Syntaxin/SNAP25 we spotted a possible third difference, a small second connection between the complex and the bilayer (Supplementary Fig. 5B-C, green arrows). These differences likely represent structural changes due to the presence of SNAP25 and support our observations of the formation of an acceptor Munc18/Syntaxin/SNAP25 complex.

**Figure 9.**
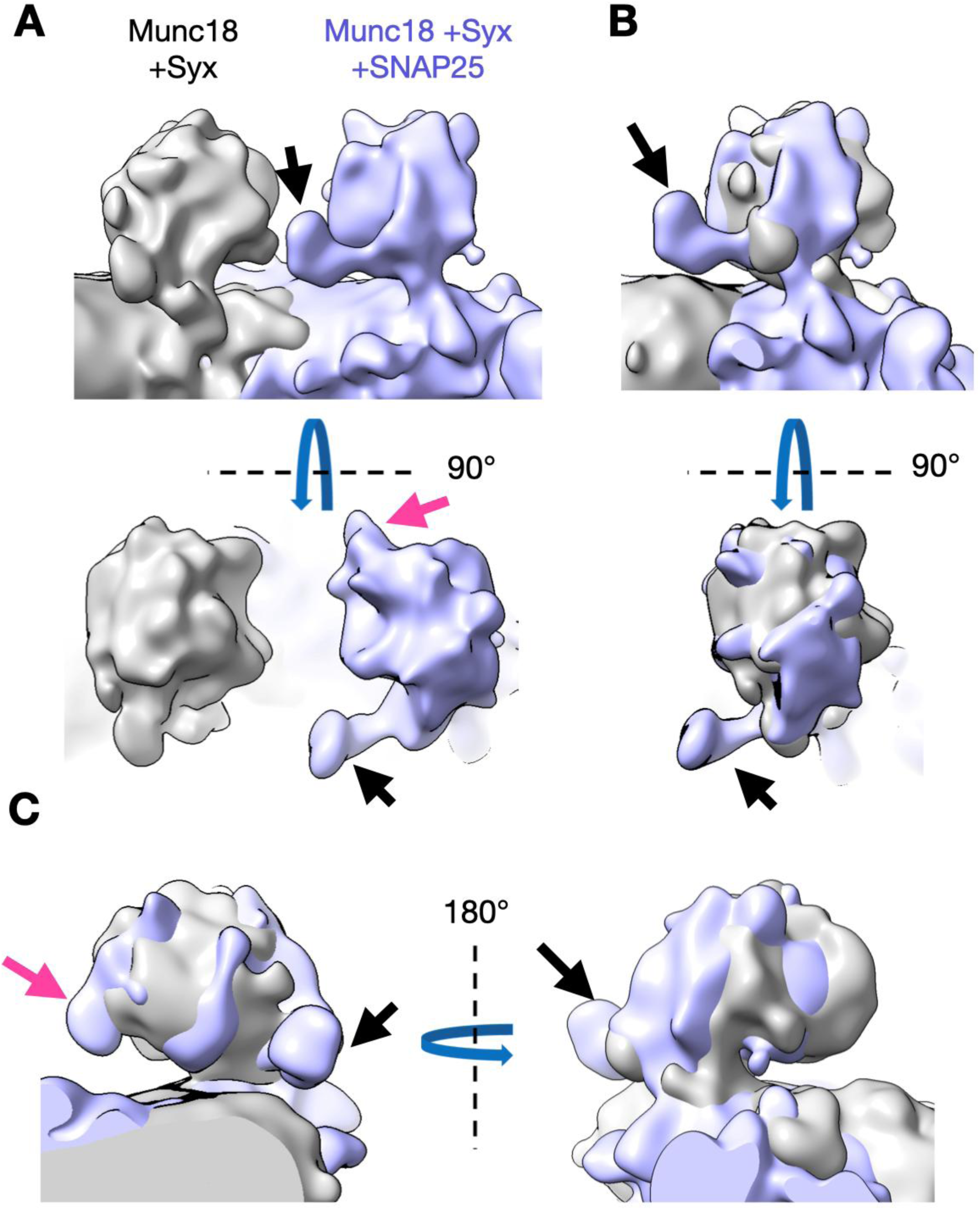
Comparison of the protein densities for the 3D reconstructions of Munc18/Syntaxin and Munc18/Syntaxin/SNAP25 in nanodiscs. **(A)** Side-by-side comparisons of the two density maps when opened together in ChimeraX. **(B)** Comparison of the protein density in the two maps when they are fit into each other in ChimeraX. **(C)** Additional views of the comparison of the two maps when they are fit into each other in ChimeraX. The black and pink arrows point to the main and secondary extra densities, respectively, that are observed in Munc18/Syntaxin/SNAP25 but lacking in Munc18/Syntaxin.

While we can observe differences in the structures, the relatively low resolution of our maps does not allow us to reliably model a precise protein arrangement in the complexes. We attempted to dock higher resolution PDB models from previous studies [31, 55] into the density maps. The crystal structure of soluble Syntaxin in complex with Munc18 (PDB 3C98) docked well, albeit not perfectly, into the Munc18/Syntaxin nanodisc structure (Supplementary Fig. 5A). Docking the same model (PDB 3C98) into the Munc18/Syntaxin/SNAP25 nanodisc structure resulted in a much poorer fit (Supplementary Fig. 5B). At this resolution, the additional density flanking the core complex is consistent with the Habc domain of Syntaxin, with the structural model fitting the observed density reasonably well (Supplementary Fig. 5C). We roughly docked the H_abc_ domain into this region of the structure and then used part of the model of the cryo-EM structure of Munc18, Syntaxin, and Synaptobrevin (PDB 7UDB) [31] to model how the SNARE domain of Syntaxin and Munc18 could be positioned in this structure (Supplementary Fig. 5C). A cartoon explaining this interpretation, given the low resolution admittedly with some liberties, is shown in Supplementary Fig. 5D.

Alternatively, it is also feasible that the extra density might correspond to SNAP25 fully or partially engaged with Syntaxin. This scenario is more difficult to model since SNAP25 is known to be largely unstructured outside of the SNARE complex. It is also known that the two SNARE motifs of SNAP25 fold and assemble with Syntaxin in a sequential pathway with the N-terminal motif inserting into the complex first while the C-terminal motif is still unfolded [9, 17]. Although this could explain the flexibility of the additional density that we observed, it would be virtually impossible to model this into the current structure. We therefore favor the interpretation given in Supplementary Fig. 5.

It is important to note that while we can approximate the positions of the proteins in our maps using existing PDB models, the fits are not perfect, and higher resolution structures or better dynamical models are needed to visualize the precise arrangements of the proteins in the complex. For example, when we mapped distances measured in the FLIC experiment (Fig. 5) into the cryo-EM map of the Munc18/Syntaxin/SNAP25 complex in proteoliposomes, we could see that the distances measured by FLIC fall within the protein density (Supplementary Fig. 6). However, due to the low resolution we cannot confidently model any protein arrangement in this map. The structures obtained from nanodisc samples have slightly higher resolution, but fitting FLIC distances in those maps is challenging due to the high flexibility of the complex relative to the nanodisc (Fig. 8B and E).

## Discussion

In this study we show that the lipid environment and presence of SNAP25 regulate Syntaxin’s lateral organization in model membranes and that these factors modulate and in turn get modulated by the binding of the SNARE-assembly chaperone Munc18. We have used two methods to measure the oligomerization of Syntaxin in membranes. They assay different aspects of the oligomerization likely at different length scales and in different regions of the protein. The SDS resistance assay extracts Syntaxin from its membranes, disperses it, and records remaining interactions, presumably all occurring through the SNARE motifs and very little through the transmembrane domains. Small scale oligomers consisting of just a few molecules of Syntaxin and SNAP25, if present, remain and are measured in these assays. By contrast, the fluorescence self-quenching assay records interactions between an unspecified number of Syntaxins while they are in a lipid bilayer environment. This assay does not distinguish between small oligomers and large clusters of oligomers, but it records the oligomerization state of proteins in their natural lipid bilayer environment that is not perturbed by detergents.

Syntaxin is more oligomerized and clustered in bPC:Chol than in PM 1% PIP2 membranes, but it gets slightly dispersed by SNAP25 in both environments. However, when extracted with SDS from both membranes, SDS-resistant dimers of Syntaxin persist. When SNAP25 is also present, higher order SDS-resistant homo- and hetero-oligomers with stoichiometries 1:1, 2:1, and 2:2 are observed independent of the membrane background (Fig. 1).

Munc18 binds best to Syntaxin/SNAP25 complexes in PM 1% PIP2 membranes (Fig. 2). The absence of SNAP25 reduces Munc18 binding to Syntaxin, and changing the lipids to bPC:Chol, i.e. increasing clustering of Syntaxin or Syntaxin/SNAP25, decreases Munc18 binding to Syntaxin or Syntaxin/SNAP25 complexes. We also observed a modest effect of Munc18 on Syntaxin’s clustering in model membranes, with opposite effects in PM 1% PIP2 and bPC:Chol membranes. The observed interplay of Munc18- and lipid-dependent Syntaxin clustering in model membranes is in agreement with previous studies in cells showing that Syntaxin’s interactions with surrounding lipids are crucial for regulating its lateral organization in the plasma membrane [16, 23]. Our observation that Munc18 slightly promotes formation of SDS-resistant Syntaxin/SNAP25 complexes also supports the idea of the formation of a ternary acceptor SNARE complex consisting of Munc18, Syntaxin and SNAP25 [28]. That previous study also showed that ternary Munc18/Syntaxin/SNAP25 complexes are not very stable which is consistent with our co-floatation results with proteoliposomes (Fig. 3). In PM 1% PIP2 membranes containing co-reconstituted Syntaxin/SNAP25, some Munc18 remains attached and co-floats with the Syntaxin/SNAP25 complex, but a sizable fraction comes off the complex during centrifugation. The Munc18/Syntaxin complex (without SNAP25) appears to be more stable in this assay.

Our FLIC microscopy results further show that Munc18 can form a complex with Syntaxin alone and Syntaxin/SNAP25 and elevate these complexes including Syntaxin’s H_abc_ domain from the membrane surface. This action of Munc18 prevents the SNARE motif(s) from interacting with the lipid bilayer surface [9, 17, 41], makes them more accessible to interact with the SNARE motif of Synaptobrevin, and allows them to engage with other regulatory proteins including Munc13, Complexin, and Synaptotagmin.

The lipid-dependent oligomerization of Syntaxin has an impact on the docking and fusion of insulin granules with planar supported membranes. Docking is more efficient in PM 1% PIP2 compared to bPC:Chol membranes, and the fusion kinetics are faster, likely due to higher accessibility of Syntaxin in PM 1% PIP2. We show that Munc18 interacts more efficiently and in a different mode with Syntaxin/SNAP25 in PM 1% PIP2 compared to bPC:Chol membranes (Fig. 2 and 3). Because of these binding differences, we observe that in PM 1% PIP2 membranes the addition of Munc18 slightly reduces the number of docking-competent sites, increases fusion probability, and slows the kinetics of fusion, suggesting that fusion is mediated by Munc18/Syntaxin/SNAP25 complexes. In bPC:Chol membranes however, Munc18 significantly reduces the number of docking-competent sites but does not affect the fusion kinetics, leading to the conclusion that fusion is mediated by Syntaxin/SNAP25 complexes. Therefore, it is likely that Munc18 controls the correct assembly of the SNARE complex [30, 31, 44–50] more efficiently in PM 1% PIP2 than in bPC:Chol membranes.

We have shown previously that calcium-triggered fusion of insulin granules is kinetically delayed in the absence of Munc13 due to low levels of CAPS associated with purified insulin granules, which resulted in incomplete priming of these granules [53]. It is possible that the kinetic delay that we observed here in PM 1% PIP2 membranes in presence of Munc18, but in the absence of Munc13 and calcium indicates that additional assembly steps of SNAREs must occur in order to fully prime secretory vesicles for efficient calcium-triggered membrane fusion. Together, these experiments lay the groundwork for future studies to dissect the membrane-embedded molecular mechanisms governing priming and calcium-triggered fusion and to define the roles of key regulatory proteins, including Munc13, complexin, and synaptotagmin.

The interactions of Munc18 with Syntaxin alone and the Syntaxin/SNAP25 Q-SNARE complex that we observed on reconstituted membranes prompted us to pursue a cryoEM structure of the lipid and membrane bound complexes. Even though the complexes were somewhat labile and likely flexible, we were able to image enough complexes in nanodiscs and proteoliposomes to observe differences between the Munc18/Syntaxin and Munc18/Syntaxin/SNAP25 structures. This is significant since, except for the post-fusion full *cis*-SNARE complex [4], there are, to date, few structural studies of Q-SNARE and Munc18/SNARE complexes in membranous environments.

Our two cryo-EM structures in nanodiscs suggest that the protein arrangement is quite different between the Munc18/Syntaxin and Munc18/Syntaxin/SNAP25 structures. The most striking feature is that a domain that we tentatively attribute to the H_abc_ domain of Syntaxin is dislocated to one side to make room for SNAP25 in the Munc18/Syntaxin/SNAP25 structure compared to the Munc18/Syntaxin structure. The latter seems to resemble a previously determined soluble Munc18/Syntaxin structure [55], while the former would be consistent with a study that suggested that Syntaxin’s conformation in Munc18/Syntaxin/SNAP25 complex is more open than in the Munc18/Syntaxin complex [28]. In the Munc18/Syntaxin/SNAP25 structure we also observed a thin density (pink arrows in Fig. 9) on the opposite side of the complex from the neck density connecting the head of the complex with the nanodisc. This could be part of SNAP25 pointing towards the nanodisc or it could represent a conformational shift in Munc18. A third feature (green arrows in Supplementary Fig. 5B and C) that we observed at low density thresholds only in the Munc18/Syntaxin/SNAP25 structure and not in the Munc18/Syntaxin structure is a second connection of the protein density to the density of the nanodisc. A feature visible only at low-density thresholds likely reflects partial occupancy, whether due to structural flexibility or substoichiometric incorporation. It is plausible that this density belongs to SNAP25 in the region where it is anchored in the bilayer. Likewise, it seems reasonable to attribute the main density of the neck connecting the globular protein head with the nanodisc that is present in both structures to the linker between the SNARE motif and transmembrane domain of Syntaxin. Due to the limited resolution of our structures, we were unable to build a detailed model of the exact arrangements and conformations of each protein in these two complexes. Nevertheless, we do detect reproducible key differences between them.

The structures that we observed are very likely stoichiometric structures containing single copies of each protein in both structures. Given the tendency of SNAREs to oligomerize as observed in this study, it is possible, perhaps even likely that complexes of additional stoichiometries exist in our samples (Supplementary Fig. 3). However, such additional assemblies might contain strong conformational variability preventing proper classification by the employed standard processing pipelines.

We made extensive efforts to improve the resolution of our structures. For both nanodisc structures we tried to improve the resolution by approximately tripling the amount of data for each condition. Increasing the data sample size did not improve the resolution. Although several membrane protein structures have been solved with predominantly side views [56, 57], we also considered the possibility that an insufficient number of different views could be responsible for the low resolution of our structure (Supplementary Fig. 7). Inspection of projection images in cryoSPARC (Supplementary Fig. 8A) revealed a strong preference for side views with top or bottom views being largely absent from the particle population (red boxes in Supplementary Fig. 8A). While particles in these orientations were identifiable in the micrographs (Supplementary Fig. 8B), none of our picking and classification strategies reliably selected or retained them. To facilitate future analyses with more advanced picking algorithms, the smaller and larger datasets used in this study have been deposited in EMPIAR (see Supplementary Table 1).

Structural studies of SNARE proteins are technically challenging because SNAREs are small with large unstructured regions except in the fully formed *cis*-SNARE complex [9, 17, 41]. They can also form heterogenous complexes even within carefully prepared samples [5–9]. There have been attempts to gain structural insights into SNARE complex assembly intermediates using soluble protein fragments and, in some cases, using chemical crosslinking to stabilize complexes [3, 31, 55]. However, as this work demonstrates, the membrane environment affects protein-protein interactions between SNAREs themselves and between SNAREs and regulatory proteins. Despite these challenges, our previous experience with cryoEM of very small and heterogenous complexes of other proteins [58–60] makes us confident that solving high-resolution structures of SNAREs and their complexes with other proteins within membrane environments could be feasible in the future. Adding additional regulatory proteins (e.g. Munc13, complexin) could increase the uniformity and size of the particles and thereby may facilitate structural studies of such complexes.

### Experimental procedures

#### Materials

The following materials were purchased and used without further purification: porcine brain L-α-phosphatidylcholine (bPC), porcine brain L-α-phosphatidylethanolamine (bPE), porcine brain L-α-phosphatidylserine (bPS), L-α-phosphatidylinositol (liver, bovine) (PI), phosphatidylinositol 4,5-bisphosphate [PI(4,5)P_2_], 1,2-dioleoyl-snglycero-3-phosphoethanolamine-N-(lissamine rhodamine B sulfonyl) (Rh-DOPE), and 1,2-dioleoyl-sn-glycero-3-phosphoethanolamine-N(7-nitro-2-1,3-benzoxadiazol-4-yl) (NBD-DOPE) were from Avanti Research (Alabaster, AL); DMPE-PEG2000triethoxysilan (DPS) was custom-synthesized by Shearwater Polymers (Huntsville, AL); cholesterol, sodium cholate, EDTA, potassium chloride, calcium (CaCl_2_), OptiPrep Density Gradient Medium, sucrose, MOPS, potassium acetate were from Sigma; L-glutamic acid potassium salt monohydrate and SDS were from Research Products International; HEPES was from Bio Basic; ovalbumin was from GE Healthcare; Nycodenz, glycerol, chloroform, ethanol, Contrad detergent, all inorganic acids and bases, and hydrogen peroxide were from Fisher Scientific. Water was purified first with deionizing and organic-free 3 filters (Virginia Water Systems) and then with a NANOpure system from Barnstead to achieve a resistivity of 18.2 megohms/cm.

Alexa Fluor dyes were from Thermo Fisher; Janelia Fluor 552 was kindly gifted by the Janelia Research Campus. The Hex18A peptide for nanodisc preparation was custom-synthesized by GenScript. Lacey Carbon 300 Copper mesh grids were purchased from Ted Pella, Inc. (catalog number 01895). Antibodies for Syntaxin-1a (mouse monoclonal, catalog number 110 011), SNAP-25 (rabbit polyclonal, catalog number 111 002), and Munc18 (mouse monoclonal, catalog number 116 011) were from Synaptic Systems.

#### Protein purification and labeling

Syntaxin-1a, SNAP-25 and Munc18 from *Rattus norvegicus* were expressed under T7 promoter in pET28s expression vector in *Escherichia coli* strain BL21(DE3) cells. Proteins were purified as described before [34, 52] starting with Ni-NTA affinity chromatography. N-terminal His-tag was removed by thrombin cleavage and the proteins were further purified using ion-exchange and size-exclusion chromatography when necessary. SNAP25 palmitoylation was mimicked by dodecylation of its four native cysteines with dodecyl methanethiosulfonate (Toronto Research Company) [34]. We refer to dodecylated SNAP25 as dSNAP25.

Syntaxin-1a (residues 1-288) used in the experiments had no native cysteines present (C271A, C272A). Syntaxin used for the fluorescence quenching assay had an additional cysteine at the C-terminus labeled with Alexa Fluor 647 maleimide (Thermo Fisher) [10, 11]. Syntaxin constructs used for sdFLIC experiments have a single cysteine at the residue of interest labeled with Alexa Fluor 546 maleimide (Thermo Fisher). His-tagged syntaxin-1a was reacted with at least two-fold molar excess of Alexa-546 in thoroughly degassed DPC-buffers. Labeled proteins were separated from free dye via extensive wash after re-binding to a Ni-NTA column. Subsequently, eluted Alexa-labeled proteins were subjected to thrombin cleavage and then purified by size-exclusion chromatography [41].

Lysine residues in Munc18 were labeled with Alexa Fluor 546 NHS ester (Thermo Fisher). A five-fold molar excess of Alexa-564 NHS ester was reacted with the protein for 2h at room temperature to achieve labeling of 3 lysines per protein molecule on average. Unbound dye was removed using PD10 columns (Cytiva).

#### SNARE protein reconstitution in proteoliposomes and planar supported membranes

Lipids were mixed in ratios bPC:Chol 70:30 or bPC:bPE:bPS:Chol:PI:PI(4,5)P_2_, 25:25:15:30:4:1. Organic solvents were evaporated by N_2_ gas and further dried by vacuum desiccation for at least 1h. Lipid films were then dissolved in 25 mM sodium cholate in a buffer (20 mM HEPES, 150 mM KCl, pH 7.4). Syntaxin-1a and dSNAP25 were added to achieve a 3,000:1; 1,000:1 or 400:1 lipid:protein ratio depending on the experiment, and the mixture was allowed to equilibrate at room temperature for 1h. Samples were then dialyzed against 5 liters of buffer (20 mM HEPES, 150 mM KCl, pH 7.4) overnight at 4°C with a buffer change after ∼4h.

Planar supported membranes with reconstituted Q-SNAREs were prepared by the Langmuir-Blodgett/vesicle fusion technique as described previously [61, 62]. Quartz slides were cleaned using Piranha solution (3:1 sulfuric acid: hydrogen peroxide) and rinsed with MilliQ water. The first leaflet of the bilayer was deposited on the slide using Langmuir-Blodgett transfer using a Nima 611 Langmuir-Blodgett through. In all experiments, a composition of bPC:Chol:DPS (70:30:3) was used for the monolayers. The lipid mixture in chloroform was deposited on the water surface in the trough, solvent was allowed to evaporate, and the monolayer was compressed to reach a target pressure of 32 mN/m^2^. The slides were quickly dipped then slowly withdrawn with a computer maintaining constant surface pressure. The second leaflet of the bilayer was formed by incubating monolayer-covered slides inside a perfusion chamber with Q-SNARE containing proteoliposomes for 1h at room temperature. Excess proteoliposomes were removed by perfusing the chamber with 10 ml of buffer (20 mM HEPES, 150 mM KCl for binding assay; or 20 mM HEPES, 120 mM potassium glutamate, 20 mM potassium acetate for the single granule fusion assay, both pH 7.4).

#### Florescence self-quenching assay

Syntaxin-1a (1-289C) labeled at the C-terminus with Alexa Fluor 647 maleimide (Thermo Fisher) was reconstituted in proteoliposomes alone or with dSNAP-25 at protein:lipid ratio 1:1,000 (each protein) and Syntaxin:dSNAP25 ratio of 1:1 when dSNAP25 was present. Proteoliposomes were diluted 75 times in buffer (20 mM HEPES, 150 mM KCl, pH 7.4) and the fluorescence intensity was measured at room temperature before and after 15 min incubation with 0.5 µM Munc18 on Fluorolog-3 spectrofluorimeter (HORIBA Jobin Yvon). Excitation was at 650 nm and an emission scan between 660 and 700 nm with 1 nm increment was collected. 4 nm slits were used for both excitation and emission. Integrated intensities of the fluorescent peaks between 660-682 nm were used to quantify the fluorescence intensities.

#### SDS-resistant SNARE oligomers

Formation of SDS-resistant homo-oligomers of Syntaxin and hetero-oligomers of Syntaxin and dSNAP25 was investigated as described in Margittai et al. [37]. Syntaxin was reconstituted alone or co-reconstituted with dSNAP25 in proteoliposomes at a protein:lipid ratio 1:1,000, each protein. Samples were incubated with 19 µM Munc18 (to achieve the same Munc18 excess over Syntaxin as used in fluorescence self-quenching assay) for 15 min at room temperature, then SDS was added without boiling and samples were separated by SDS-PAGE in 4-20% Mini-PROTEAN TGX gels (BioRad).

#### Western blots

Samples from the Mini-PROTEAN TGX gels (BioRad) were transferred to nitrocellulose membranes (BioRad), blocked with 5% milk in Tris-buffered saline with 0.1% Tween 20 (TBST), and analyzed by Western blotting with anti-Syntaxin and anti-SNAP25 antibodies. All primary antibodies were used at a 1:1,000 dilution in 5% milk TBST. Secondary antibodies conjugated with IRDye 680 or IRDye 800 were used at 1:10,000 dilution in 5% milk TBST. Blots were imaged using Odyssey Imaging System (LI-COR).

#### Protein binding to planar supported membranes

Imaging chambers with planar supported membranes containing reconstituted SNARE proteins (at protein:lipid ratio 1:3,000) were placed on a Total Internal Reflection Fluorescence (TIRF) microscope (for details, see below). An image of the field of view was taken every 30 seconds and the average intensity of that region was recorded. The bilayers were imaged for 2.5 min to collect the background signal before the fluorescently labeled Munc18 (0.5 µM) was injected into the imaging chamber, and the binding curve was collected. When ovalbumin was used to block unspecific Munc18-bilayer interaction, the planar supported membranes were incubated with 2 µM ovalbumin for 15 min, then 0.5 µM Munc18 was added without washing out the ovalbumin.

#### Co-floatation assay

Syntaxin and dSNAP25 were co-reconstituted in proteoliposomes at a protein:lipid ratio 1:3,000, each protein. Proteoliposomes were incubated with 0.5 µM Munc18 for 15 minutes at room temperature before being loaded onto the Nycodenz gradient in polycarbonate centrifuge tubes (Beckman Coulter, catalog number 343775). 50 µl of 80% Nycodenz + 50 µl of proteoliposomes (40% final Nycodenz concentration) were mixed at the bottom of the tube, 50 µl of 30% Nycodenz was layered on top of that, followed by 50 µl of buffer (20 mM HEPES, 150 mM KCl, pH 7.4). Samples were then centrifuged at 197,000 x g for 1.5h using TLS-55 rotor. Ten 20 µl fractions were collected from top to bottom, loaded onto a Mini-PROTEAN TGX gels (BioRad), and analyzed by Western blotting against Munc18 and Syntaxin.

#### sdFLIC microscopy

The principle of site-directed fluorescence interference contrast (sdFLIC) microscopy and the set up used in this work has been described previously [41, 43]. A membrane containing Syntaxin with specifically labeled cysteines (at protein:lipid ratio 1:3,000) is supported on a patterned silicon chip with microscopic steps of silicon dioxide. The fluorescence intensity depends on the position of the dye with respect to the standing modes of the exciting and emitting light in front of the reflecting silicon surface. The position is determined by the variable-height 16 oxide steps and the constant average distance between dye and silicon oxide [63].

Images were acquired on a Zeiss Axiovert 200 or Axio Observer 7 fluorescence microscope (Carl Zeiss) with a mercury lamp as a light source and a 40× water immersion objective (Zeiss; N.A. = 0.7). Fluorescence was observed through a 610-nm band-pass filter (D610/60; Chroma) by a CCD camera (DV-887ESC-BV; Andor-Technologies). Exposure times for imaging were set between 40 and 80 ms, and the excitation light was filtered by a neutral density filter (ND 1.0, Chroma) to avoid photobleaching.

During sdFLIC experiments, we acquired 4-6 images, 20-30 min after buffer changes, for each membrane condition of one supported membrane. From each image, we extracted 100 sets of 16 fluorescence intensities and fitted the optical theory with the fluorophore-membrane distance as fit parameter. Software to fit the data was kindly provided by the authors of [63]. The standard deviation of these ∼400-600 results were usually on the order of 1 nm. The optical model consists of 5 layers of different thickness and refractive indices (bulk silicon, variable silicon oxide, 4 nm water, 4 nm membrane, bulk water), which we kept constant for all conditions [41, 42, 64]. The reported errors for the absolute membrane distance are the standard errors from at least 3 repeats. Not included in these errors are systematic errors that might originate in different membrane thicknesses or membrane-substrate distances between different lipid conditions and a systematic underestimation of the residue-membrane distance from 10-20% of protein that is trapped on the substrate proximal side of the supported bilayer. The reported errors after the addition of Munc18 are the standard errors of the detected distance changes from at least 3 repeats for each condition. Based on previous experiments with polymer supported bilayers we estimate the systematic uncertainty for the measured absolute distance to be ∼ ±1 nm [64].

#### Sample preparation for cryo-EM

Proteoliposomes were prepared at a final lipid concentration of 2 mM and a protein:lipid ratio 1:1,000. Samples were incubated with 10 μM Munc18, i.e. a 5-fold molar excess of Munc18 over Syntaxin for 15 min at room temperature and either directly frozen on the grid or used to generate nanodiscs. For nanodisc preparation, 1,900 µl of each sample was concentrated to 700 µl (loading volume for size- exclusion chromatography) and incubated with Hex18A peptide (amphipathic peptide modified with hexanoic acid of the N-terminus) [54] at a final concentration of 4 mg/ml for 2h on ice with gentle mixing every 20-30 min. The samples were then purified on a Superdex200 10/300 GL column to remove free peptide. 0.5 ml fractions were collected and fractions from each elution peak were combined (fractions from each peak were combined separately), concentrated and immediately frozen on the EM grids. Lacey 300 Cu Mesh carbon grids were glow discharged for 20 seconds with a current of 25 mA using a GLOQUBE plus^TM^. To make the surface properties of the grids more favorable for our sample a piece of filter paper with several drops of amylamine was added to the glow discharge chamber with the grids. This results in the grids being relatively hydrophobic and positively charged. Vitrobot was used to flash-freeze the samples, 3.2 µl of the sample was loaded per grid, blotting was done with force set to 5 for 5 sec.

#### Cryo-EM single particle reconstruction

All cryo-EM imaging was performed at the University of Virginia Molecular Electron Microscopy core. The general workflows for cryo-EM reconstruction for the conditions are shown in Supplementary Fig. 9. The cryo-EM imaging and reconstruction parameters are shown in Supplementary Table 1. For all protein complexes many different image processing pipelines were tried. Standard automating particle picking procedures in cryoSPARC such as “Blob Picking” or “Template Picking” were unable to reliably pick the protein density on the surface of the nanodiscs or proteoliposomes. Therefore, for each condition’s dataset, several thousand particles were manually picked. These manual particles were picked so that the center of each particle was the apparent protein density on the surface of the bilayer (Supplementary Fig. 9). It should be noted that protein densities on the surface of the proteoliposomes or nanodiscs were quite heterogenous. This is exemplified by looking at the densities on the surface of the proteoliposome shown in Supplementary Fig. 3.

For Munc18/Syntaxin/SNAP25 in proteoliposomes manually picked coordinates were then used as direct inputs into Topaz [65]. For both nanodisc conditions 2D classification was performed on the manually picked particles and particles from selected 2D classes were used as inputs for Topaz. For nanodisc conditions, even with Topaz-picked particles, protein density showed up in the 2D class averages only if (1) the “Re-center 2D classes” parameter was turned off or (2) if the re-centering was kept on but the “Re-center mask threshold” was increased to very high values (0.9995 used) from the default value of 0.2. Similar procedures were found to be necessary with the proteoliposome dataset.

For all datasets the Topaz-picked particles were extracted from micrographs and sorted using 2D classification. Particles from selected classes were then input into ab initio reconstruction jobs. For the nanodisc conditions only a single class was used for ab initio reconstruction while for the proteoliposomes two classes were used to sort out additional heterogeneity. For ab initio reconstructions, the volumes and particles from ab initio reconstruction were input into homogenous refinement with a low pass filter of 30 Å applied to each volume. The resolutions of the structures in this paper are reported in Supplementary Table 1. “Gold standard” Fourier shell correlation curves are shown in Supplementary Fig. 7. For all structures we were unable to achieve high enough resolution to see secondary structure features such as α-helices. A variety of masking procedures were tried to reach higher resolution, but none of them were successful. Our conclusion is that sample heterogeneity (flexibility, possibly different multimeric states, etc.) coupled with the relatively small size of the protein complexes and lack of symmetry makes high resolution structure determination difficult under the current sample conditions.

UCSF ChimeraX [66] was used for analyzing the density maps and for docking previously published models into them.

#### INS1 and GRINCH cell culture

Rat insulinoma (INS1) cells [67] were cultured on 10-cm plastic cell culture plates in RPMI medium supplemented with 10% FBS, 10 mM HEPES, 1 mM sodium pyruvate, 50 mM β-mercaptoethanol, 1x antibiotic-antimycotic at 37°C and 5% CO_2_. GRINCH cells (INS1 cells expressing C-peptide-GFP, [68]) were cultured as described above, with the addition of 20 µg/ml of G418 (Gibco) selection in the media. GRINCH cells were additionally transiently transfected with VAMP2-Halo plasmid via electroporation using ECM 830 Electro Square Porator (BTX). Cells were collected from the plates and suspended in a small volume of cytomix electroporation buffer (120 mM KCl, 10 mM KH_2_PO_4_, 0.15 mM CaCl_2_, 2 mM EGTA, 25 mM HEPES, 5 mM MgCl2, 2 mM ATP, 5 mM glutathione, pH 7.6) with the addition of 30 μg DNA per electroporation cuvette. Two 8 ms pulses at 255 V were applied. Cells were then transferred to regular 10 cm culture plates with regular growth medium and cultured for 3 days before being harvested for isolation of secretory granules. However, the VAMP2-Halo tag was not utilized in the current work.

#### Insulin granule purification from INS1 and GRINCH cells

Insulin granules were purified from 20-30 plates of INS1 or GRINCH cells following a previously described protocol [51, 53]. Cells were scraped into PBS and pelleted via centrifugation, then washed by resuspension in the homogenization buffer (0.26 M sucrose, 5 mM MOPS, 0.2 mM EDTA) and pelleted again. The cells were resuspended in the homogenization buffer with the addition of protease inhibitor cocktail (Thermo Scientific) and cracked open with a ball bearing homogenizer (0.2507 inch bore and 0.2496 diameter ball). The homogenate was subjected to two rounds of centrifugation in a fixed-angle micro-centrifuge: first at 1,500 x g for 10 min to remove nuclei and large debris, then at 11,200 x g for 15 min to remove mitochondria. The post-mitochondrial supernatant was collected and adjusted to 5 mM EDTA. If granules were isolated from cells transfected with VAMP2-Halo, Janelia Fluor 552 (1 mM final concentration in the sample) was added and incubated for 2h at room temperature. The sample was then layered on a discontinuous Optiprep (iodixanol) gradient in Beckmann SW55 tubes: 30% iodixanol at the bottom (0.5 ml), 14.5% iodixanol in the middle (3.3 ml), and post-mitochondrial supernatant at the top (1.2 ml). The Optiprep solutions were prepared by first mixing 60% Optiprep with buffer (0.26 M sucrose, 30 mM MOPS, 1 mM EDTA + protease inhibitor) in 5:1 ratio to make 50% iodixanol working solution and then diluting it further with homogenization buffer. The sample was spun at 190,000 x g for 5h. A white band at the interface of the 30% and 14.5% Optiprep gradient was collected as the insulin granules. The sample was dialyzed into fusion assay buffer (120 mM potassium glutamate, 20 mM potassium acetate, 20 mM HEPES, pH 7.4) for 48-72h at 4°C (three 5 liter changes of buffer). Granules were then flash frozen in 10% glycerol and stored at -80°C.

#### FRET-based bulk fusion assay

Proteoliposomes with Syntaxin and dSNAP25 (protein:lipid ratio 1:400) were prepared so that they also contained 1.5% each of the fluorescent lipid probes NBD-DOPE and Rhodamine-DOPE that form a fluorescence resonance energy transfer (FRET) pair. Proteoliposomes were diluted 100-fold in the reaction buffer (120 mM potassium glutamate, 20 mM potassium acetate, 20 mM HEPES, pH 7.4) and incubated at 37°C for 15 min with the addition of 0.5 μM Munc18 when used. NBD fluorescence was recorded over time on Fluorolog-3 spectrofluorimeter (HORIBA Jobin Yvon), with excitation at 460 nm and emission at 532 nm and 1 sec time increment using 4 nm slits in the excitation and emission paths. After baseline fluorescence was collected for 5 min, 100 μl of insulin granules isolated from INS1 cells were added, an increase of NBD fluorescence due to FRET relief was observed for 20 min, and finally 0.1% Triton was added to record the maximum NBD fluorescence in the fully solubilized sample, which was taken as the value of 100% fusion to normalize each experiment.

#### Single granule fusion assay

Planar supported membranes with Syntaxin and dSNAP25 at a protein lipid ratio 1:3,000 (each protein) were formed on a quartz slide as described above. Bilayers were washed with 10 ml of fusion buffer (120 mM potassium glutamate, 20 mM potassium acetate, 20 mM HEPES, pH 7.4) with addition of 100 µM EDTA. When Munc18 was included, bilayers were additionally perfused with 2 ml of 0.5 µM Munc18 in fusion buffer and incubated for 15 min. 50 µl of secretory granules were then diluted into 1 ml of fusion buffer (also including 0.5 µM Munc18 when relevant) and injected into the chambers with the planar supported membranes. Fluorescence of the C-peptide-GFP fluorescence was immediately recorded on a TIRF microscope with a 488 nm laser and using a 200 ms exposure time.

#### TIRF microscopy

Experiments examining Munc18 binding to planar supported membranes as well as single-granule docking and fusion were imaged using Zeiss AxioObserver Z1 fluorescence microscope, with a 63x water immersion objective and a prism-based TIRF illumination. The light source was an OBIS 561 laser (Coherent Inc.) for the Munc18 binding assay and OBIS 488 laser (Coherent Inc.) for the single granule fusion assay. An evanescent wave that decays exponentially from the quartz-water interface with a penetration depth of ∼100 nm was achieved by totally internally reflecting a laser beam at an angle of 72° from the surface normal. Fluorescence was recorded by an electron-multiplying charge-coupled device (EMCCD, DV887ECS-EV, Andor Technology) that was cooled to -70°C. The laser intensity, shutter, and camera were controlled by a program written in-house in LabView (National Instruments).

#### Data analysis

Munc18 binding curves were fitted with an exponential offset function in IgorPro (Wavemetrics), following the equation:

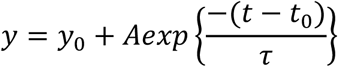

Single granule TIRF images were analyzed using a homemade Matlab (Mathworks) program to detect docked granules and extract the peak and mean fluorescence intensities of 5 px x 5px regions of interest. A custom-made program written in LabView (National Instruments) was then used to categorize events and obtain data about docking-fusion delay times for each fusion event as described previously [69].

Cumulative distribution functions (CDFs) of the fusion delay times were generated using GraphPad Prism software. The CDFs were then fitted in IgorPro with a parallel reaction model according to the equation [52, 69]:

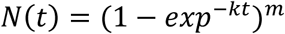

where t is time, k is a rate constant, and m is the number of parallel reactions occurring. The characteristic time to reach ½ of max fusion probability T_1/2max_ was calculated from fitted values for k and m. The reported errors for T_1/2max_ were determined through error propagation of the fit errors.

GraphPad Prism software was used to create plots and statistical analysis of the data. For statistical analysis of two experimental variables, t-tests were used. Modifications to account for non-Gaussian distribution of the data and unequal SDs were applied when relevant. For statistical analysis of more than two experimental variables, ANOVA was used, with the correction for non-Gaussian distribution of the data applied when relevant.

## Data availability

CryoEM datasets were deposited with EMPIAR. CryoEM structures were deposited in the EMDB as EMD-76033, EMD-76034, and EMD-76039 respectively for Munc18/Syntaxin in nanodiscs, Munc18/Syntaxin/SNAP25 in nanodiscs, and the Munc18/Syntaxin/SNAP25 in proteoliposomes.

## Supporting information

This article contains Supporting Information.

## Acknowledgements

We would like to thank Klaudia Lukow for help with culturing GRINCH cells and the purification of insulin granules. We thank Prisha Shah for assistance with bulk proteoliposome fusion experiments. We thank Dr. Binyong Liang for purification of the proteins used in the FLIC experiment. We thank Dr. Edward Egelman for use of his cryo-EM workstations as well as some methodological advice regarding cryo-EM analysis. We thank Dr. Ravi Sonani for assistance EMPIAR depositions. Lastly, we would like to thank Dr. Michael Purdy and Dr. David Cooper at the University of Virginia Molecular Electron Microscopy Core for their assistance with cryo-EM screening and data collection.

## Funding and additional information

This work was supported by NIH grant P01 GM072694 and institutional funds.

