## supplementary information for "Munc18 binds to and organizes membrane-bound acceptor Q-SNARE complexes in a fashion that depends on the membrane’s lipid composition"

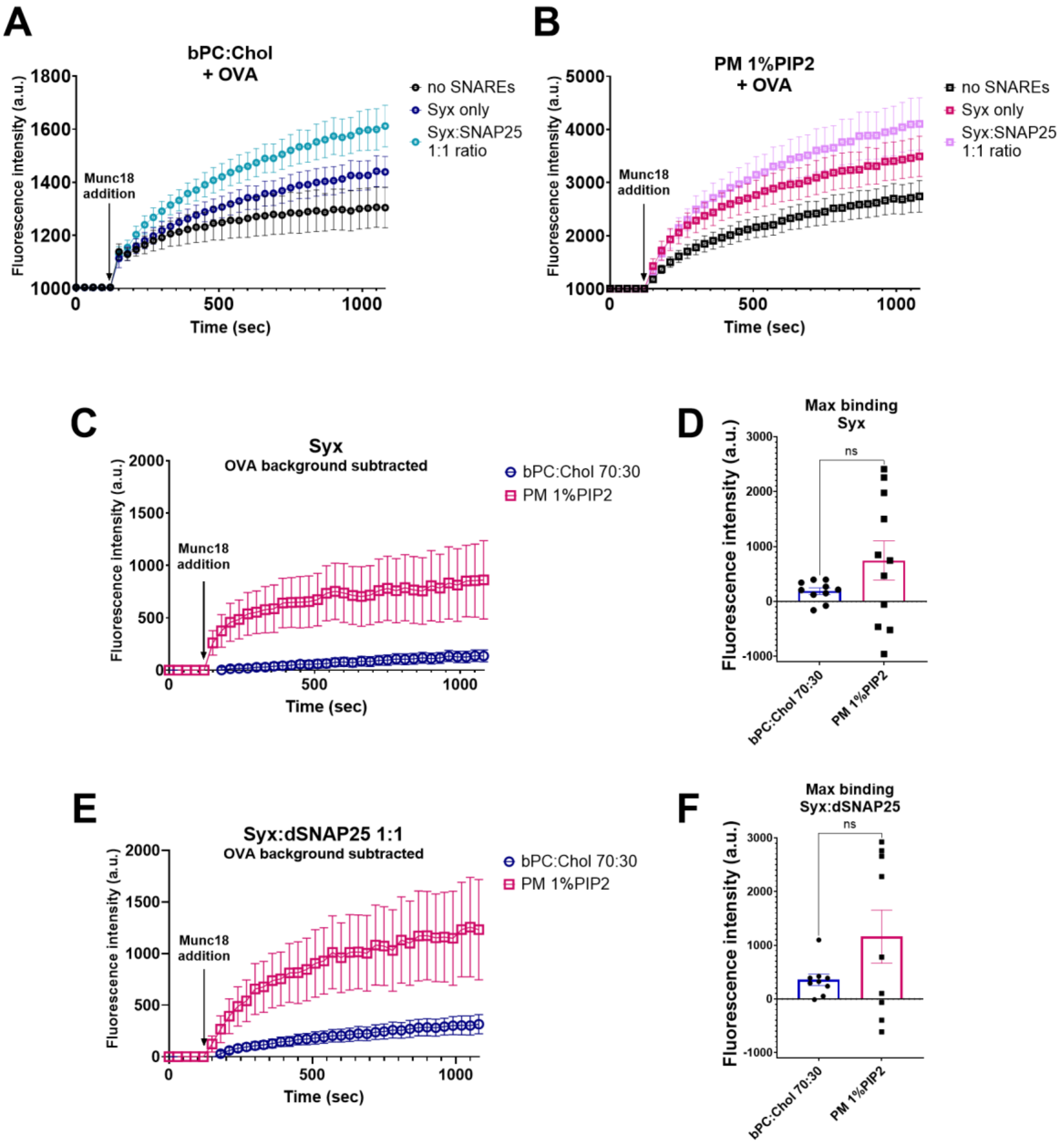


**Supplementary Figure 1. Ovalbumin was used to block unspecific Munc18-membrane interactions in planar supported membranes. (A and B)** TIRF microscopy binding curves of 0.5 µM Munc18 to bPC:Chol 70:30 **(A)** and PM 1% PIP2 **(B)** planar supported membranes in the presence of ovalbumin. **(C)** TIRF microscopy binding curves of 0.5 µM Munc18 to Syntaxin only bilayers with pure lipid and ovalbumin (no SNAREs) background subtracted. **(D)** Maximum fluorescence intensity values obtained from fits of the binding curves from **(C)** to 1$^{st}$ order kinetic functions. **(E)** TIRF microscopy binding curves of 0.5 µM Munc18 to Syntaxin/SNAP25 bilayers with pure lipid and ovalbumin (no SNAREs) background subtracted. **(F)** Maximum fluorescence intensity values obtained from fits of the binding curves from **(E)** to 1$^{st}$ order kinetic functions. Error bars represent standard error. T-test with Welch's correction for unequal SDs was used in **(D)** and **(F)**.

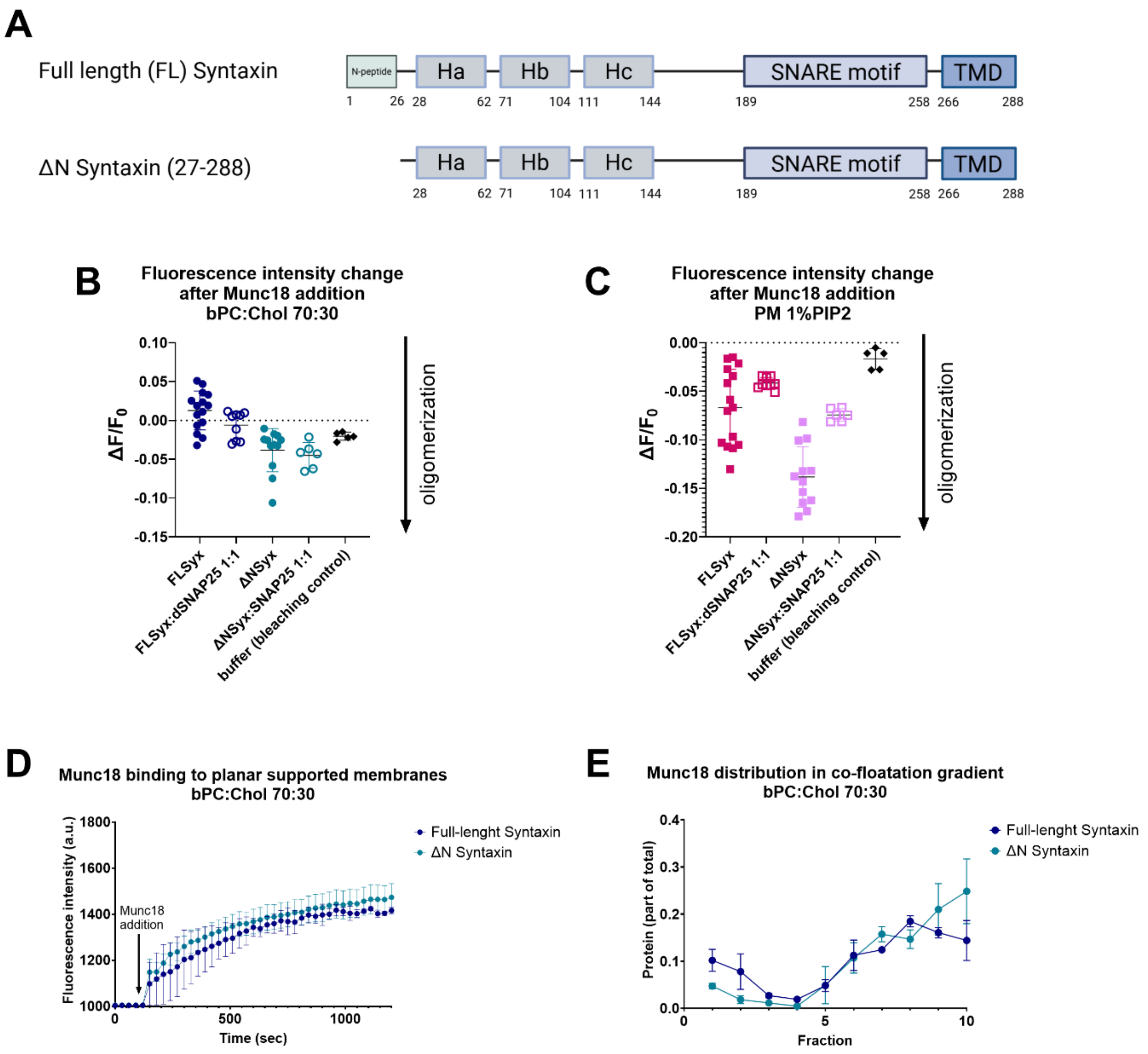


**Supplementary Figure 2. Syntaxin's N-peptide affects Munc18 effect on Syntaxin's oligomerization, but not the Munc18-Syntaxin interaction. (A)** Domain structures of Syntaxin with and without the N-peptide (residues 1-26). **(B and C)** Effect of addition of 0.5 µM Munc18 on full-length Syntaxin's and ΔN Syntaxin's oligomerization probed by Alexa 647 fluorescence quenching in bPC:Chol **(B)** and PM 1% PIP2 **(C)** bilayers. **(D)** Binding of 0.5 µM Munc18 to Syntaxin and ΔN Syntaxin in planar supported bPC:Chol membranes. **(E)** Distribution of Munc18 in co-floatation assay with full-length Syntaxin and ΔN Syntaxin proteoliposomes.

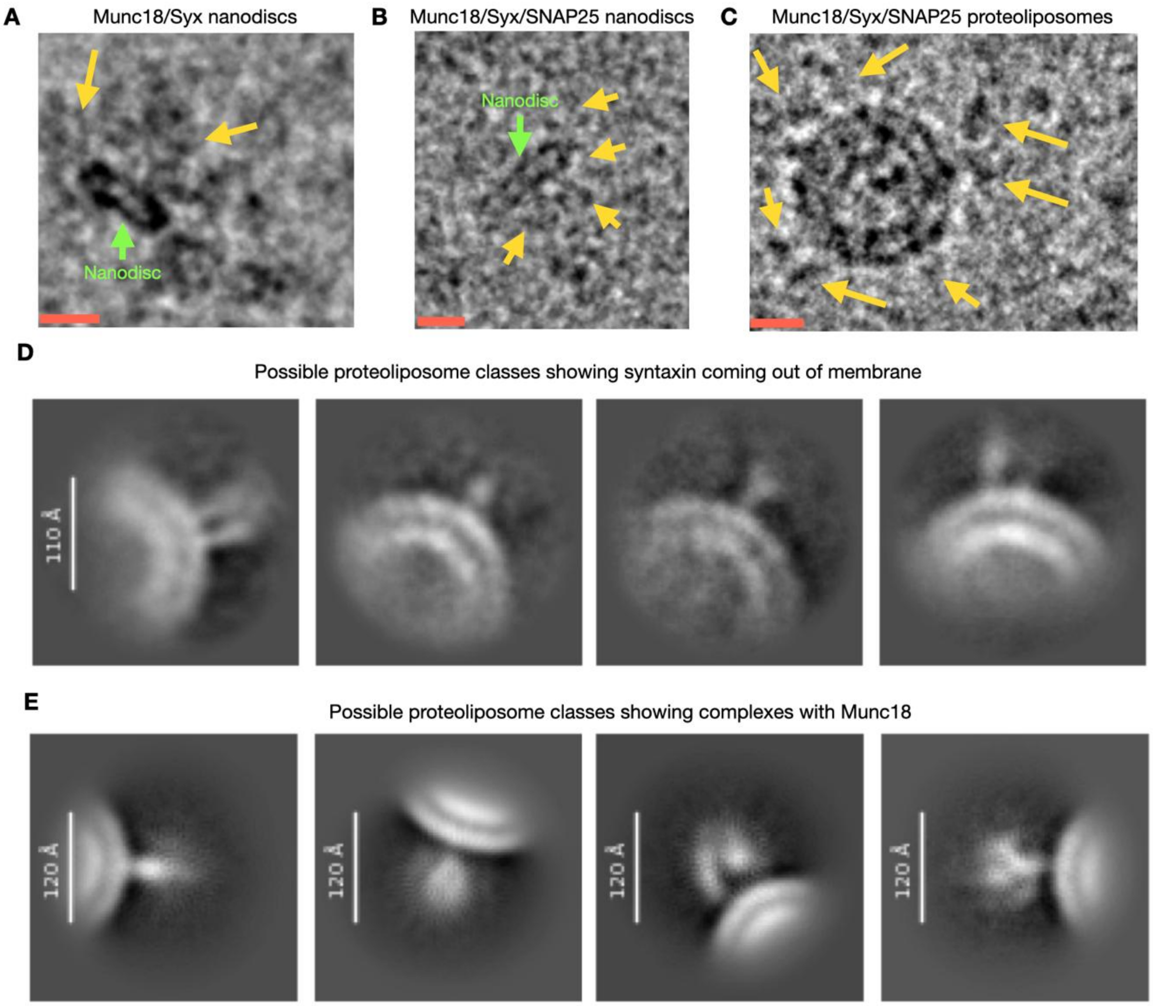


**Supplementary Figure 3. Examples of heterogeneity in protein density on the surface of (A) Munc18/Syntaxin in nanodiscs, (B) Munc18/Syntaxin/SNAP25 in nanodiscs, and (C) Munc18/Syntaxin/SNAP25 in proteoliposomes.** The yellow arrows point to protein density on the particle surface. The scale bar is ~100 Å. Note that the sizes of the various densities on the particle surface are variable, which could indicate heterogeneity in the protein assemblies on the surface. **(D)** 2D class averages from the Munc18/Syntaxin/SNAP25 in proteoliposomes showing what could be just Syntaxin projecting from the membrane. **(E)** 2D class averages from the Munc18/Syntaxin/SNAP25 in proteoliposomes showing what appears to be a protein complex with a larger globular protein bound to the complex on the surface of the membrane.

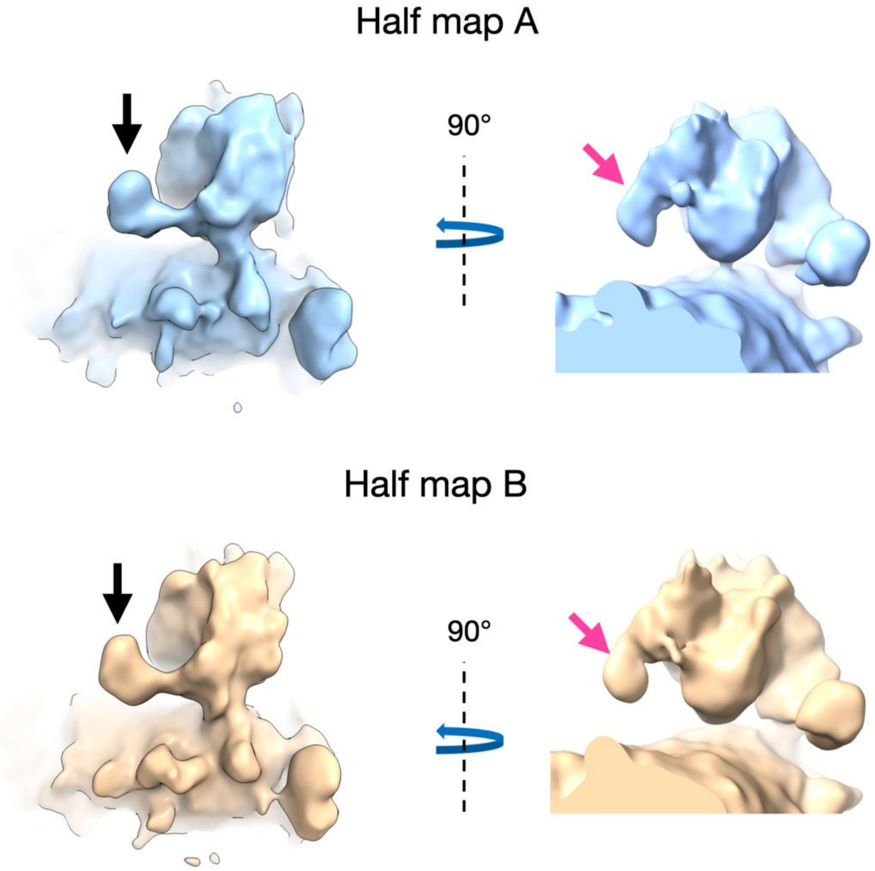


**Supplementary Figure 4**. **Half maps for the Munc18/Syntaxin/SNAP25 cryo-EM reconstruction.** The arrows point to the main and secondary densities of interest, also described in Figure 9, extending from the structures in both half maps. For visualization, the half maps were filtered to the appropriate resolution of comparison (~13 Å) in ChimeraX.

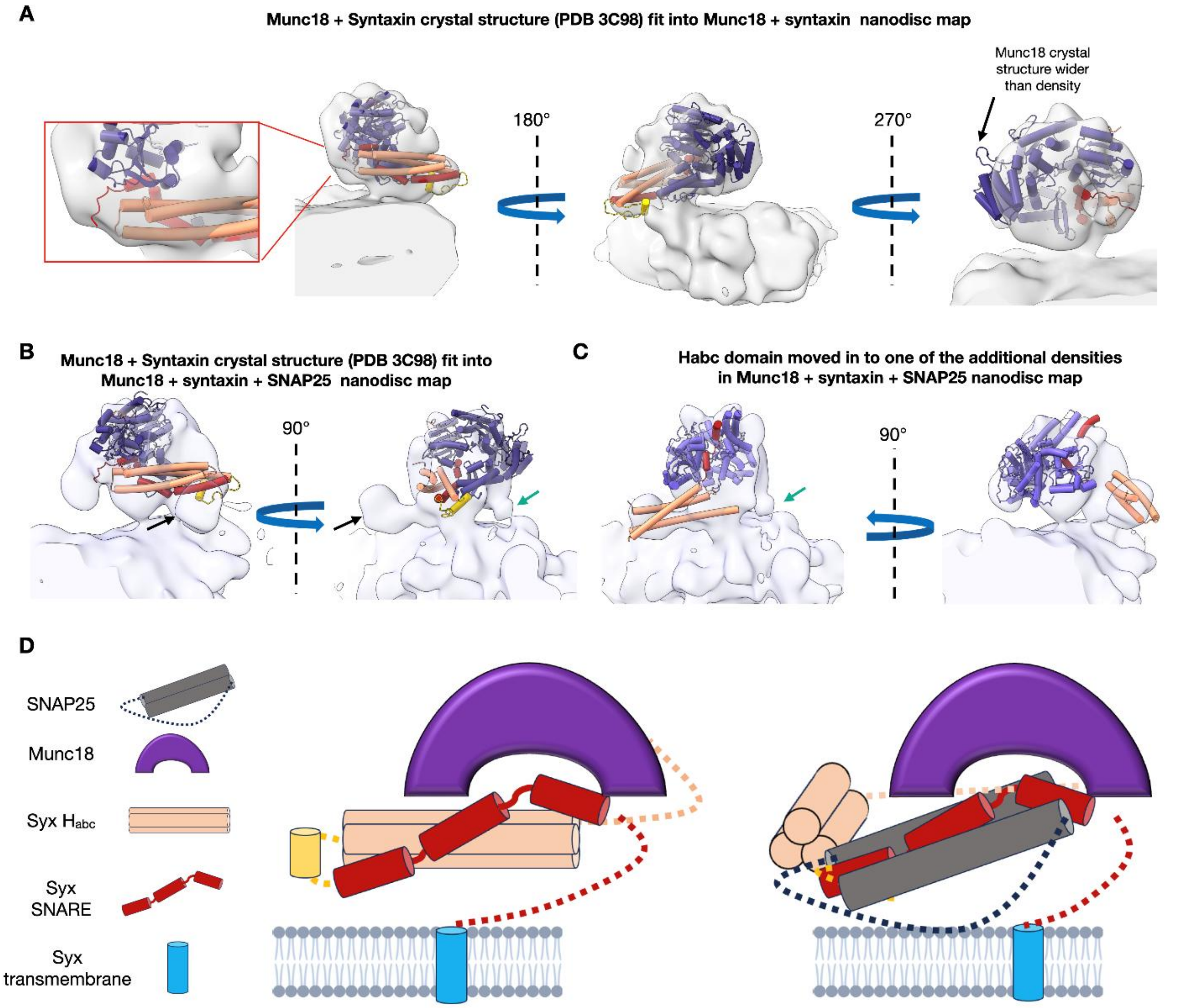


**Supplemental Figure 5. PDB models of higher resolution component structures can be used to interpret the nanodisc cryo-EM density maps.** Munc18 is colored purple in all panels. The $H_{abc}$ domain, SNARE motif, and linker between those domains of Syntaxin are colored light pink, red, and gold respectively. **(A)** Docking of the crystal structure of Munc18 and Syntaxin (PDB 3C98) into the Munc18/Syntaxin nanodisc density map. **(B)** The same model (PDB 3C98) is docked into the Munc18/Syntaxin/SNAP25 nanodisc density map. The black arrow points to a large density that the model cannot fit in its current conformation. The green arrow points to a small density that touches the bilayer at lower density thresholds. **(C)** In this alternate model the $H_{abc}$ domain of Syntaxin from PDB 3C98 has been moved into the large density. The SNARE motif and Munc18 from PDB 7UDB were utilized in this model to better fit the cryo-EM density map. It is possible that experimental cryo-EM density with no high-resolution structural models fit could be partially occupied by SNAP25. **(D)** Cartoon roughly interpreting our observations from these low resolution cryo-EM maps without (left) and with (right) SNAP25 (in gray) included.

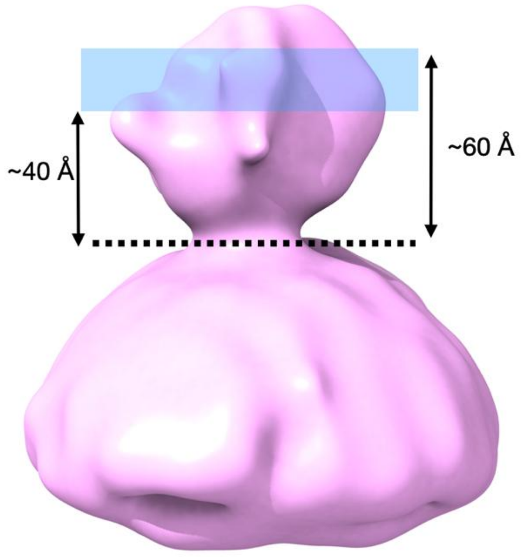


**Supplementary Figure 6. Range of distances of Syntaxin residues (blue area) probed by FLIC microscopy and mapped into the cryo-EM map of the Munc18/Syntaxin/SNAP25 complex in PM 1% PIP2 proteoliposomes.** The dotted line is the approximate location of the membrane surface.

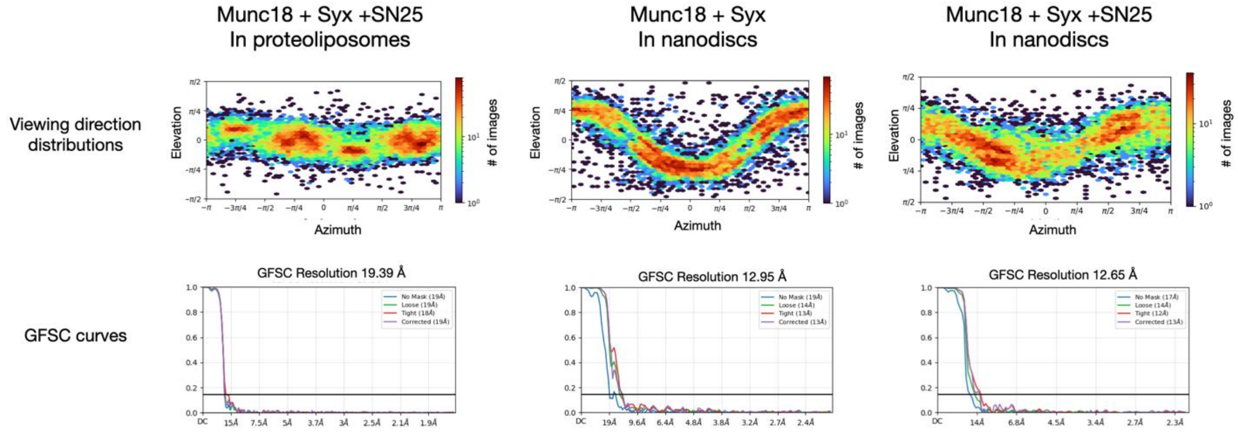


**Supplementary Figure 7. CryoSPARC orientation distributions (top) and Map:Map "Gold standard" Fourier shell correlation (GFSC) curves (bottom) for the three cryoEM structures in this study.**

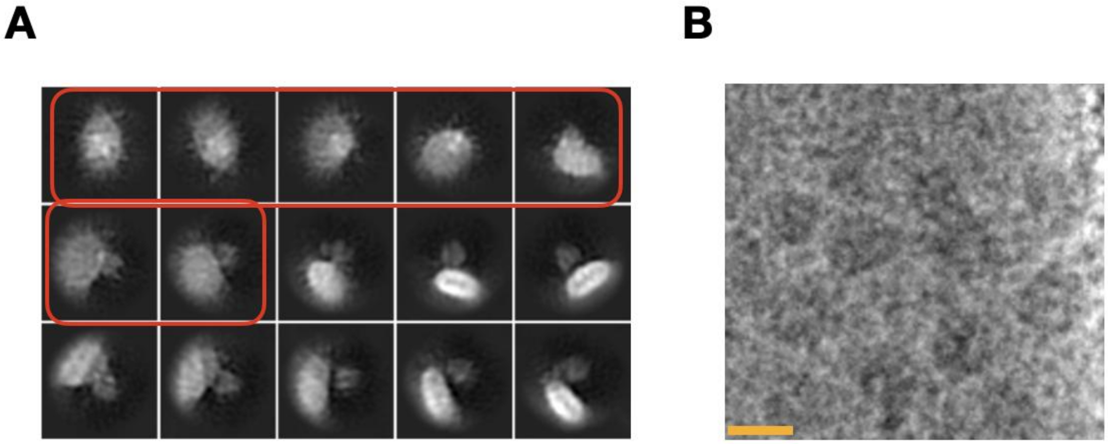


**Supplementary Figure 8. (A)** 2D projections of the Munc18/Syntaxin/SNAP25 nanodiscs created to simulate the various views for the protein complex and nanodisc. Orientations that would be considered top or bottom views have a red box around then. **(B)** Region of a micrograph containing what might be "top/bottom" views of the nanodisc, presumably with protein. Scale bar is 150 Å.

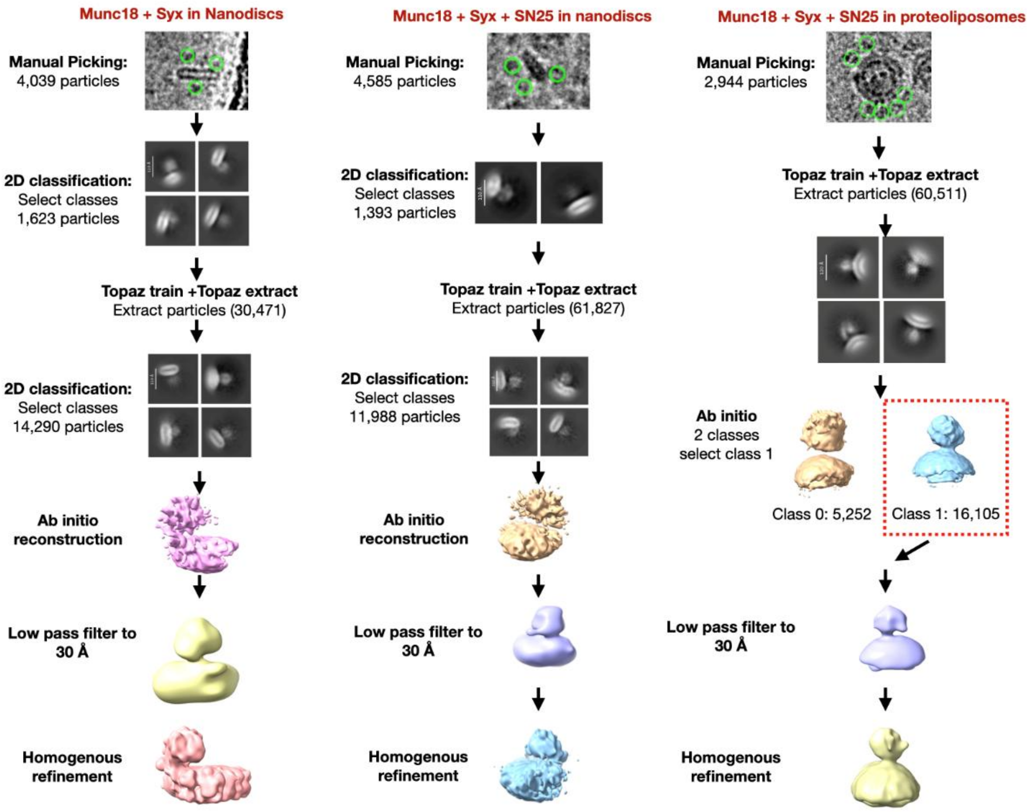


**Supplementary Figure 9. CryoSPARC workflows for single particle reconstructions.** Workflows are shown for the reconstructions of Munc18/Syntaxin in nanodiscs (left), Munc18/Syntaxin/SNAP25 in nanodiscs (middle), and Munc18/Syntaxin/SNAP25 in proteoliposomes (right).

**Supplementary Table 1. Cryo-EM reconstruction statistics**

| Munc18/Syx in Nanodiscs | |
|---|---|
| EMDB ID | EMD-76033 |
| EMPIAR deposition dataset 1 | EMPIAR-13651 |
| EMPIAR deposition second dataset | EMPIAR-13649 |
| Number of movies | 3,006 |
| Microscope | Titan Krios |
| Pixel size (Å/pixel) | 1.065 |
| Voltage (keV) | 300 |
| Spherical aberration (mm) | 2.7 |
| Total dose (e/Å^2) | 60 |
| Number of particles | 14,290 |
| 0.143 map:map FSC resolution (Å) | 13 |
| **Munc18/Syx/SN25 in Nanodiscs** | |
| EMDB ID | EMD-76034 |
| EMPIAR deposition dataset 1 | EMPIAR-13652 |
| EMPIAR deposition second dataset | EMPIAR-13703 |
| Number of movies | 2,959 |
| Microscope | Titan Krios |
| Pixel size (Å/pixel) | 1.065 |
| Voltage (keV) | 300 |
| Spherical aberration (mm) | 2.7 |
| Total dose (e/Å^2) | 60 |
| Number of particles | 11,988 |
| 0.143 map:map resolution FSC (Å) | 12.7 |
| **Munc18/Syx/SN25 in proteoliposomes** | |
| EMDB ID | EMD-76039 |
| EMPIAR deposition | EMPIAR-13650 |
| Number of movies | 8,093 |
| Microscope | Titan Krios |
| Pixel size (Å/pixel) | 0.83 |
| Voltage (keV) | 300 |
| Spherical aberration (mm) | 2.7 |
| Total dose (e/Å^2) | 60 |
| Number of particles | 16,105 |
| 0.143 map:map FSC resolution (Å) | 19.4 |
